# A tissue-resolved endothelial surface proteome atlas informs organ-selective vascular targeting

**DOI:** 10.64898/2026.08.21.746320

**Authors:** Yanru Deng, Huiqiao Li, Jieyi Meng, Andrew Lemoff, Hong Zhou, Xinchun Pi, Yi Zhu

**Affiliations:** USDA/ARS Children’s Nutrition Research Center, Department of Pediatrics, Baylor College of Medicine, Houston, TX 77030, USA; Department of Biochemistry, The University of Texas Southwestern Medical Center at Dallas, Dallas, TX 75390, USA; Cardiovascular Research Institute, Baylor College of Medicine, Houston, TX 77030, USA

**Keywords:** endothelial cells, *in vivo* proximity labeling, cell surface proteomics, vascular heterogeneity, targeted drug delivery

## Abstract

**BACKGROUND:** Endothelial cells (ECs) exhibit organ-specific functional diversity that shapes vascular homeostasis, disease susceptibility, and therapeutic accessibility. Although single-cell transcriptomic studies have defined endothelial heterogeneity at the RNA level, the *in vivo* cell-surface protein landscape that directly mediates vascular signaling and drug targeting remains incompletely characterized. Here, we mapped organ-specific endothelial surface proteomes atlas *in vivo* to define tissue-enriched vascular protein candidates relevant to organ-selective therapeutic design.

**METHODS:** We generated *Cdh5*-CreERT; Cre-iPEEL mice, referred to here as CHRP mice, in which membrane-tethered horseradish peroxidase is induced selectively in ECs after tamoxifen treatment, and compared CHRP labeling with non-selective NHS-Biotin vascular labeling. Following *in vivo* biotin-phenol perfusion, endothelial surface proteins were enriched by streptavidin affinity purification and analyzed by mass spectrometry across six organs. Proteomic profiles were used to resolve tissue- and subtype-associated endothelial surface signatures, compare protein and transcript detection patterns, and nominate tissue-selective endothelial membrane candidates, which were annotated using ChEMBL compound-target information.

**RESULTS:** Compared with non-selective NHS-Biotin labeling, CHRP improved endothelial specificity and produced clearer separation of tissue-resolved endothelial surface proteomes across brain, white adipose tissue, small intestine, kidney, lung, and skeletal muscle. CHRP proteomics revealed pronounced organ-specific heterogeneity and resolved canonical arterial, venous, and capillary programs, as well as specialized endothelial signatures including blood-brain barrier and glomerular endothelial features. Comparison with single-cell endothelial references revealed systematic differences between transcriptomic and proteomic detection of endothelial membrane proteins. Further analysis identified tissue-selective endothelial membrane candidates, and ChEMBL annotation linked a subset of these candidates to existing compound-target records, supporting the candidate atlas as a resource for future tissue-selective vascular targeting studies.

**CONCLUSIONS:** CHRP-based *in vivo* proximity labeling enables systematic, protein-level mapping of organ-specific endothelial surface proteomes. Together with transcriptomic comparison and compound-target annotation, this study provides a tissue-resolved endothelial surfaceome resource for vascular biology and future organ-selective therapeutic target evaluation.

**Clinical Perspective:** *What Is New?:* CHRP-based in *vivo* proximity labeling enables systematic mapping of endothelial surface proteomes across multiple organs. The resulting atlas reveals organ-specific endothelial surface heterogeneity, including tissue-enriched metabolic, transport, receptor, and signaling signatures. Comparison with endothelial single-cell transcriptomic references identifies class-specific differences between RNA- and protein-based detection, while ChEMBL database links a subset of tissue-selective candidates to existing compound-target records.

*What Are the Clinical Implications?:* Organ-specific endothelial surface proteins provide candidates for tissue-selective vascular targeting and organ-directed therapeutic delivery. Protein-level surface mapping complements transcriptomic atlases by identifying vascular targets that may be underestimated by RNA-based approaches. The tissue-selective endothelial candidate atlas provides a resource for prioritizing both ChEMBL-annotated targets and additional surface proteins for future vascular targeting studies.

## INTRODUCTION

Endothelial cells (ECs) line the entire vascular system and exhibit remarkable functional diversity across organs, enabling precise regulation of tissue-specific vascular permeability, nutrient exchange, and immune interactions [1–3]. This heterogeneity is fundamental to both physiological homeostasis and disease susceptibility in organs such as the brain, kidney, lung, intestine, and adipose tissue [4–6]. Dysregulation of endothelial function contributes to a wide range of cardiovascular and metabolic disorders [7, 8], underscoring the importance of understanding tissue-specific endothelial specialization at the molecular level.

Recent advances in single-cell RNA sequencing have substantially advanced our understanding of endothelial diversity, identifying transcriptionally distinct endothelial subtypes across tissues and vascular beds [9, 10]. These studies have revealed arterial, venous, and capillary programs, as well as highly specialized endothelial populations such as the blood-brain barrier (BBB) and glomerular endothelium [11, 12]. However, transcriptomic profiles do not necessarily predict protein abundance, particularly for membrane-associated, secreted, and metabolically regulated proteins that mediate vascular signaling and therapeutic accessibility [13–15]. Consequently, a systematic understanding of endothelial heterogeneity at the protein level, especially *in vivo* conditions, remains limited.

Proteomic approaches have provided important insights into endothelial biology, yet most studies either rely on *in vitro* culture systems or cell isolation strategies that can disrupt native vascular architecture and microenvironmental cues [16, 17]. Genetically encoded proximity-labeling approaches such as iPEEL have recently enabled cell-type-specific surface proteomic profiling in native mouse tissues[18], but systematic *in vivo* mapping of endothelial luminal surface proteomes across multiple vascular beds remains unknown. Moreover, the EC surface-where key transporters, receptors, and therapeutic targets reside-differs substantially between *in vivo* and *in vitro* systems [19]. These limitations have hindered efforts to define organ-specific endothelial surface proteomes and to identify tissue-selective, vascular-accessible protein targets relevant to disease marker identification or engineering of tissue-specific delivery systems.

In this study, we developed and applied a genetically encoded *in vivo* proximity-labeling strategy to systematically profile endothelial surface proteomes across multiple organs. Using this approach, we mapped endothelial surface proteomic heterogeneity across six tissues, resolved vascular subtype-associated programs at the protein level, and compared proteomic and transcriptomic detection of endothelial membrane proteins. We further defined tissue-selective endothelial membrane candidates and annotated these candidates with existing compound-target information. As proof of principle for genetically directed labeling beyond ECs, we also show that the same HRP-based strategy can be redirected to adipocyte surfaces in adipose tissue, leveraging tissue-specific vascular permeability to enable labeling of parenchymal cells. Together, these studies establish an *in vivo* proteomic framework for mapping vascular cell-surface heterogeneity and for prioritizing tissue-selective membrane candidates relevant to vascular biology and future therapeutic target evaluation.

## METHODS

### Data Availability

Proteomics data used for analysis are provided in Supplementary Table S1. Code for data analyses is available at https://github.com/BCMZhulab/CHRP-proteomics. Publicly available single-cell RNA-seq references included the Murine Endothelial Cell Atlas (E-MTAB-8077) and adipose tissue datasets GSM5359340, GSM5359345, and GSM5359346. The ChEMBL 30 SQLite database was downloaded from http://ftp.ebi.ac.uk/pub/databases/chembl/ChEMBLdb/releases/chembl_30 and queried locally.

The raw mass spectrometry proteomics data have been deposited to the ProteomeXchange Consortium via the MassIVE repository with the dataset identifier MSV000102970 and ProteomeXchange accession number PXD082977.

### Animals and *in vivo* vascular surface labeling

All animal procedures were approved by the Institutional Animal Care and Use Committee of Baylor College of Medicine. Mice were maintained under specific pathogen-free conditions with *ad libitum* access to food and water. To generate endothelial HRP reporter mice, *Cdh5*-CreERT mice were crossed with Cre-iPEEL mice (JAX #037698) to produce *Cdh5*-CreERT; Cre-iPEEL offspring, referred to as CHRP mice. Tamoxifen was administered at 75 mg/kg/day for 5 consecutive days, followed by at least 7 days of washout before labeling. For nonselective vascular surface labeling, wild-type C57BL/6J mice (JAX #000664) were perfused with cell-impermeable Sulfo-NHS-LC-Biotin (NHS-Biotin; ApexBio, A8003). For cell-type-restricted proximity labeling, CHRP mice were perfused with Biotin-XX Tyramide (BxxP; ApexBio Technology, A8012), followed by hydrogen peroxide to initiate HRP-catalyzed biotinylation and antioxidant quenching to terminate labeling. Brain, epididymal white adipose tissue (eWAT), small intestine, kidney, lung, and skeletal muscle were collected for downstream analyses. For adipocyte surfaceome profiling, *Adipoq*-Cre (JAX #028020); Cre-iPEEL mice were maintained on chow or 60% high-fat diet for 12 weeks before *in vivo* labeling.

### Proteomics and computational analysis

Biotinylated proteins were enriched using streptavidin affinity purification, subjected to on-bead trypsin digestion, and analyzed by mass spectrometry. Fluorescent streptavidin staining was used to visualize *in vivo* biotinylation and assess labeling patterns. Raw mass spectrometry data were processed using Proteome Discoverer version 3.0 and searched against the UniProt mouse reference proteome. Peptide and protein identifications were filtered at 1% false discovery rate using a target-decoy strategy, and proteins identified with at least two unique peptides were retained unless otherwise indicated. Detected proteins were mapped to UniProt “true proteins” (TP), defined as proteins annotated as secreted, cell membrane, or cell surface based on UniProt Subcellular Location (CC) annotations. Label-free intensities were log□-transformed, normalized, and summarized at the tissue or replicate level as appropriate.

Downstream analyses were performed in R (version ≥4.2.0) and included dimensionality reduction, tissue correlation analysis, endothelial subtype scoring, proteomic and single-cell transcriptomic comparison, metabolic surfaceome analysis, tissue-specific candidate selection, ChEMBL compound-target annotation, Gene Ontology enrichment, and adipocyte surfaceome analysis. Detailed analytical procedures, thresholds, databases, and software packages are provided in the Supplementary Methods and Key Source.

### Statistical analysis

Statistical analyses were performed in R (version ≥4.2.0) unless otherwise specified. Biological replicates represent independent mice or independently processed tissue samples, as indicated in the figure legends. Protein abundance values were log□-transformed before quantitative comparisons. Linear modeling, nonparametric tests, Fisher’s exact tests, Gene Ontology enrichment, and pathway enrichment analyses were applied as appropriate for each dataset. Multiple-hypothesis testing was controlled using the Benjamini-Hochberg false discovery rate procedure. Correlation analyses used Pearson or Spearman coefficients, as indicated in the figure legends. Exact sample sizes, statistical tests, and significance thresholds are provided in the corresponding figure legends. No statistical methods were used to predetermine sample size, and no data were excluded unless explicitly stated.

## RESULTS

### *In vivo* vascular labeling with non-selective NHS-Biotin reveals broad but non-specific endothelial proteomes

To establish our perfusion protocol, we first used a non-selective, cell-membrane-impermeable primary amine-reactive Sulfo-NHS-biotin (NHS-Biotin) to label cell-surface proteins on the luminal surface of blood vessels *in vivo*, similar to previously described [20, 21]. NHS-Biotin is a bifunctional molecule composed of an activated NHS ester on the biotin moiety, which covalently reacts with accessible primary amines on protein surfaces, and a biotin group that enables high-affinity capture by streptavidin (Supplementary Figure 1A). Following transcardiac perfusion, tissue samples were collected, and biotinylated proteins were affinity-enriched from protein lysates using streptavidin beads and subjected to mass spectrometry analysis, with parallel histological validation to assess labeling specificity and anatomical distribution (Figure 1A).

**Figure 1.**
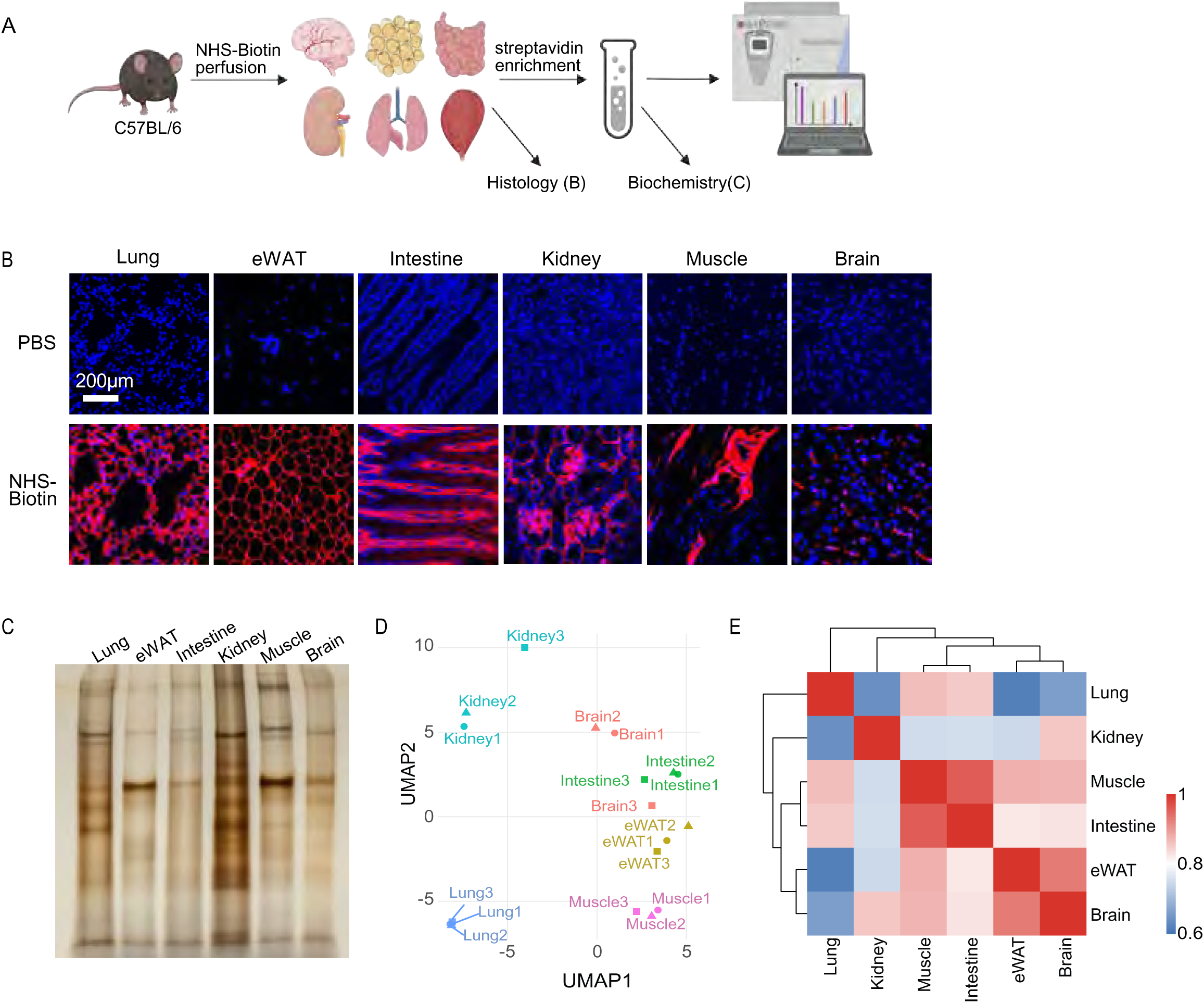
*In vivo* vascular labeling using non-selective NHS-Biotin reveals broad but non-specific endothelial-associated proteomes. **A**, Schematic illustration of the experimental workflow. C57BL/6 mice were systemically perfused with NHS-Biotin to label vascular-accessible proteins under physiological conditions, followed by tissue collection, streptavidin-based enrichment of biotinylated proteins, histological validation, and mass spectrometry-based proteomic analysis. **B**, Representative immunofluorescence images of lung, epididymal white adipose tissue (eWAT), intestine, kidney, skeletal muscle, and brain following PBS or NHS-Biotin perfusion. n=3 per group. Scale bar=200 μm. **C**, Silver staining of streptavidin-enriched protein fractions from multiple tissues after NHS-Biotin perfusion. **D**, UMAP visualization of mass spectrometry-identified proteins filtered to UniProt-annotated proteins across six tissues. **E**, Pairwise correlation analysis of tissue-averaged proteomic signatures derived from UniProt-annotated proteins following NHS-Biotin labeling. Heatmap displays Pearson correlation coefficients and hierarchical clustering between tissues.

Immunofluorescence analysis across six tissues-lung, epididymal white adipose tissue (eWAT), small intestine, kidney, skeletal muscle, and brain-revealed robust biotin labeling after NHS-Biotin perfusion (Figure 1B). However, labeling patterns varied across organs, with permeable tissues such as eWAT and small intestine showing signal beyond vascular structures, including adipocyte surfaces and intestinal villus-associated structures. Streptavidin pull-down followed by silver staining demonstrated abundant enrichment of biotinylated proteins across tissues (Figure 1C). UMAP analysis showed that lung, kidney, and skeletal muscle formed relatively distinct proteomic clusters, whereas small intestine, brain, and eWAT showed greater overlap (Figure 1D).

A similar pattern was observed when all detected proteins were examined at the individual sample level. Unsupervised clustering showed that lung and kidney samples formed relatively distinct groups, whereas brain, eWAT, small intestine, and skeletal muscle showed less complete separation (Supplementary Figure 1B). One brain replicates clustered closely with eWAT samples, and intestinal samples showed partial similarity to skeletal muscle, consistent with the overlap observed by UMAP analysis.

Pairwise correlation analysis further demonstrated high similarity among several tissues, particularly among brain, eWAT, and skeletal muscle (Figure 1E). These findings suggest that although NHS-Biotin perfusion robustly labels vascular-accessible proteins *in vivo*, its non-selective chemistry and tissue-dependent vascular permeability may introduce non-endothelial background signals that obscure tissue-specific endothelial proteomic differences.

### *In vivo* proximity labeling reveals robust tissue-specific organization of endothelial surface proteomes

The NHS-Biotin perfusion demonstrated that vascular-accessible proteins can be efficiently labeled *in vivo*, but also revealed substantial non-endothelial background signals in tissues with higher vascular permeability. We therefore sought to preserve the *in vivo* perfusion-based labeling strategy while improving endothelial specificity. The previously reported iPEEL proximity-labeling system uses membrane-tethered HRP to biotinylate extracellular proteins within the local cell-surface environment. This enzyme-dependent labeling mechanism provides genetic control over where biotinylation occurs, making it well suited for restricting surface labeling to defined cell types. We therefore reasoned that iPEEL could be adapted for endothelial-restricted surface proteomics *in vivo*. (Supplementary Figure 2A).

To achieve endothelial specificity, *Cdh5*-CreERT mice were crossed with Cre-inducible iPEEL reporter mice to generate *Cdh5*-CreERT; Cre-iPEEL (CHRP) mice. Following tamoxifen treatment, Cre-mediated recombination induced endothelial cell-specific expression of membrane-tethered HRP. Systemic perfusion with biotin-XX-phenol (BxxP), followed by streptavidin-based enrichment and mass spectrometry, enabled selective labeling and proteomic profiling of endothelial-accessible proteins (Figure 2A). Immunofluorescence analysis confirmed robust labeling across all six tissues (Figure 2B). Consistent with efficient biochemical enrichment, silver staining of streptavidin-purified fractions demonstrated substantial recovery of labeled proteins across all six tissues (Figure 2C).UMAP analysis of endothelial surface proteomes demonstrated stronger tissue-specific segregation than NHS-Biotin labeling, with biological replicates clustering tightly within each organ and separating clearly across tissues (Figure 2D). This improved organization was also evident in hierarchical clustering of all detected transmembrane and surface-associated proteins at the individual biological replicate level, which revealed coherent tissue-enriched protein blocks and grouped replicates according to tissue identity (Supplementary Figure 2B). Pairwise correlation analysis of tissue-averaged endothelial proteomic signatures further confirmed substantial divergence in endothelial protein composition across organs (Figure 2E). Together, these analyses demonstrate that endothelial-restricted CHRP-BxxP labeling improves discrimination of organ-specific endothelial surface proteomes *in vivo*.

**Figure 2.**
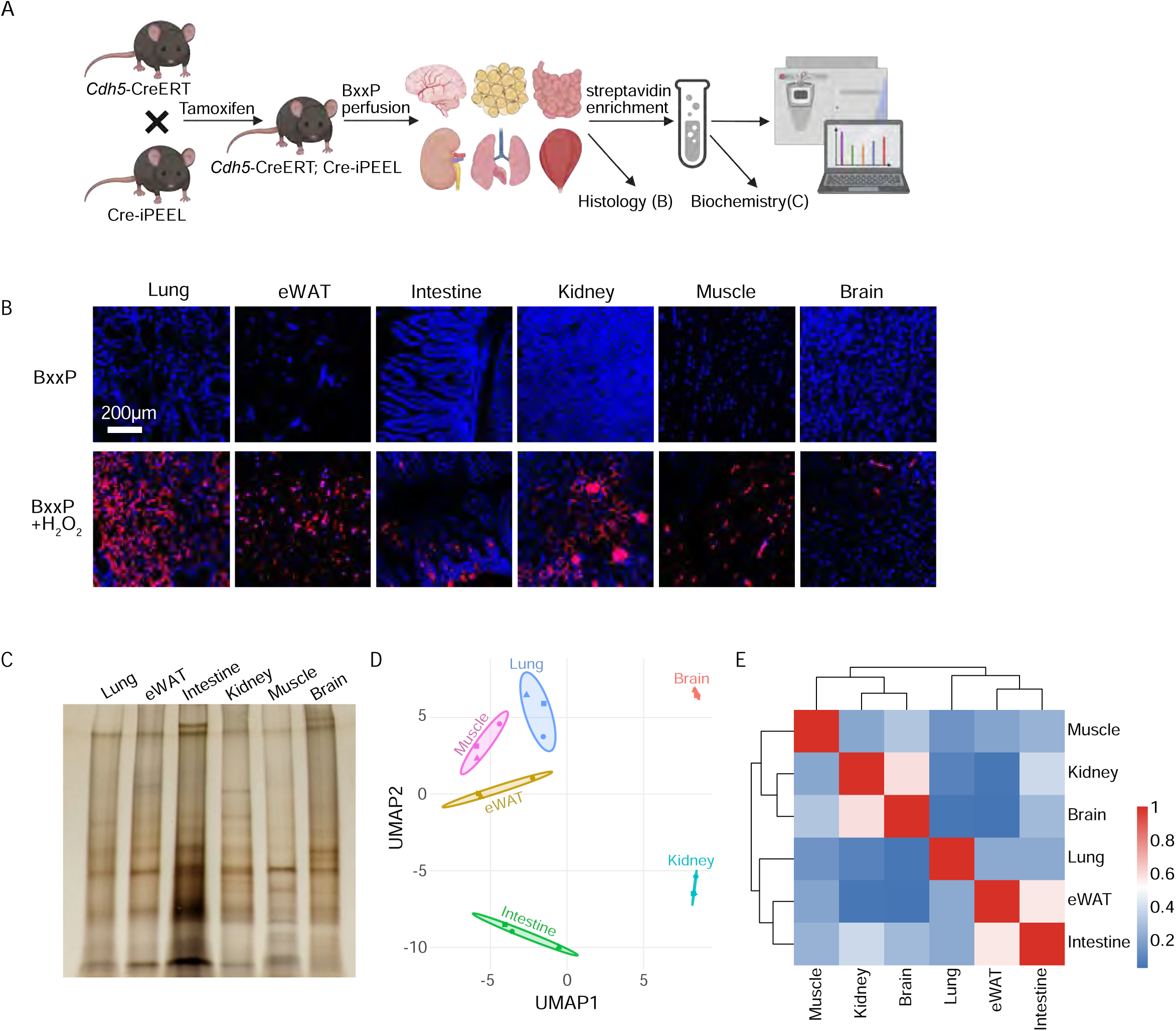
Endothelial-specific *in vivo* proximity labeling enables selective profiling of vascular surface proteomes. **A**, Schematic overview of endothelial-restricted *in vivo* proximity labeling. *Cdh5*-CreERT mice were crossed with Cre-inducible iPEEL reporter mice and treated with tamoxifen to induce endothelial-specific expression of horseradish peroxidase (HRP), referred to as CHRP mice. Following systemic perfusion with Biotin-XX Tyramide (BxxP), HRP-catalyzed proximity labeling selectively biotinylated endothelial-accessible proteins, which were subsequently enriched by streptavidin pulldown and analyzed by mass spectrometry across six tissues. **B**, Representative immunofluorescence images of lung, eWAT, intestine, kidney, muscle, and brain following BxxP perfusion. n=3 per group. Scale bar=200 μm. **C**, Silver staining of streptavidin-enriched protein fractions from indicated tissues following BxxP perfusion. **D**, UMAP visualization of UniProt-annotated proteins identified from streptavidin-enriched endothelial surface proteomes across six tissues (brain, eWAT, intestine, kidney, lung, and muscle). **E**, Pairwise correlation analysis of tissue-averaged endothelial proteomic signatures derived from UniProt-annotated proteins. Heatmap displays Pearson correlation coefficients and hierarchical clustering between tissues.

### Adipocyte-restricted HRP labeling supports cell-type-specific *in vivo* surface proteomics

To further validate the adaptability of HRP-mediated *in vivo* proximity labeling, we applied this strategy to adipocytes in adipose tissue, where high vascular permeability enables access to the adipocyte surface. Adipocyte-specific membrane-HRP mice were maintained on chow or high-fat diet (HFD), followed by BxxP perfusion, streptavidin enrichment, and mass spectrometry-based surface proteomic analysis (Supplementary Figure 3A). Immunofluorescence confirmed robust adipocyte-surface biotinylation in *Adipoq*-Cre-positive mice under both dietary conditions (Supplementary Figure 3B), and silver staining demonstrated reproducible enrichment of biotinylated proteins across biological replicates (Supplementary Figure 3C).

Unsupervised proteomic embedding separated chow- and HFD-fed adipocyte surfaceomes, indicating that adipocyte-restricted HRP labeling captured condition-dependent surface proteomic remodeling *in vivo* (Supplementary Figure 3D). Additional pathway, overlap, and differential analyses further supported diet-associated changes in the adipocyte surface proteome (Supplementary Figure 3E-G). Together, these data provide proof-of-principle evidence that genetically restricted HRP-mediated proximity labeling can be extended beyond endothelial cells to profile cell-type-associated surface proteomes *in vivo*.

### Proteomic deconvolution resolves endothelial subtype composition across vascular beds

Having established tissue-resolved endothelial surface proteomes, we next assessed whether bulk proteomic profiles retain sufficient biological structure to resolve endothelial subtypes across vascular beds. To this end, we applied three complementary deconvolution strategies using identical curated endothelial subtype marker sets (Supplementary Table S2): z-score-based marker aggregation, single-sample gene set enrichment analysis (ssGSEA) [36, 37], and non-negative least squares (NNLS) [38, 39], representing marker-based, rank-based, and linear modeling frameworks, respectively.

All approaches captured broad separation between major endothelial states; however, their ability to resolve tissue-restricted subtypes differed substantially (Figure 3A; Supplementary Figure 4A, B). Among these, z-score-based marker aggregation produced the most coherent and biologically consistent subtype landscape, recovering canonical endothelial niches, including BBB and choroid plexus endothelial cells in brain, glomerular endothelial cells in kidney, and arterial-capillary enrichment in high-flow organs such as lung and skeletal muscle. In contrast, ssGSEA exhibited reduced subtype contrast, particularly for restricted endothelial populations, whereas NNLS showed attenuated or diffused signals for specialized subtypes such as BBB and glomerular endothelial cells.

**Figure 3.**
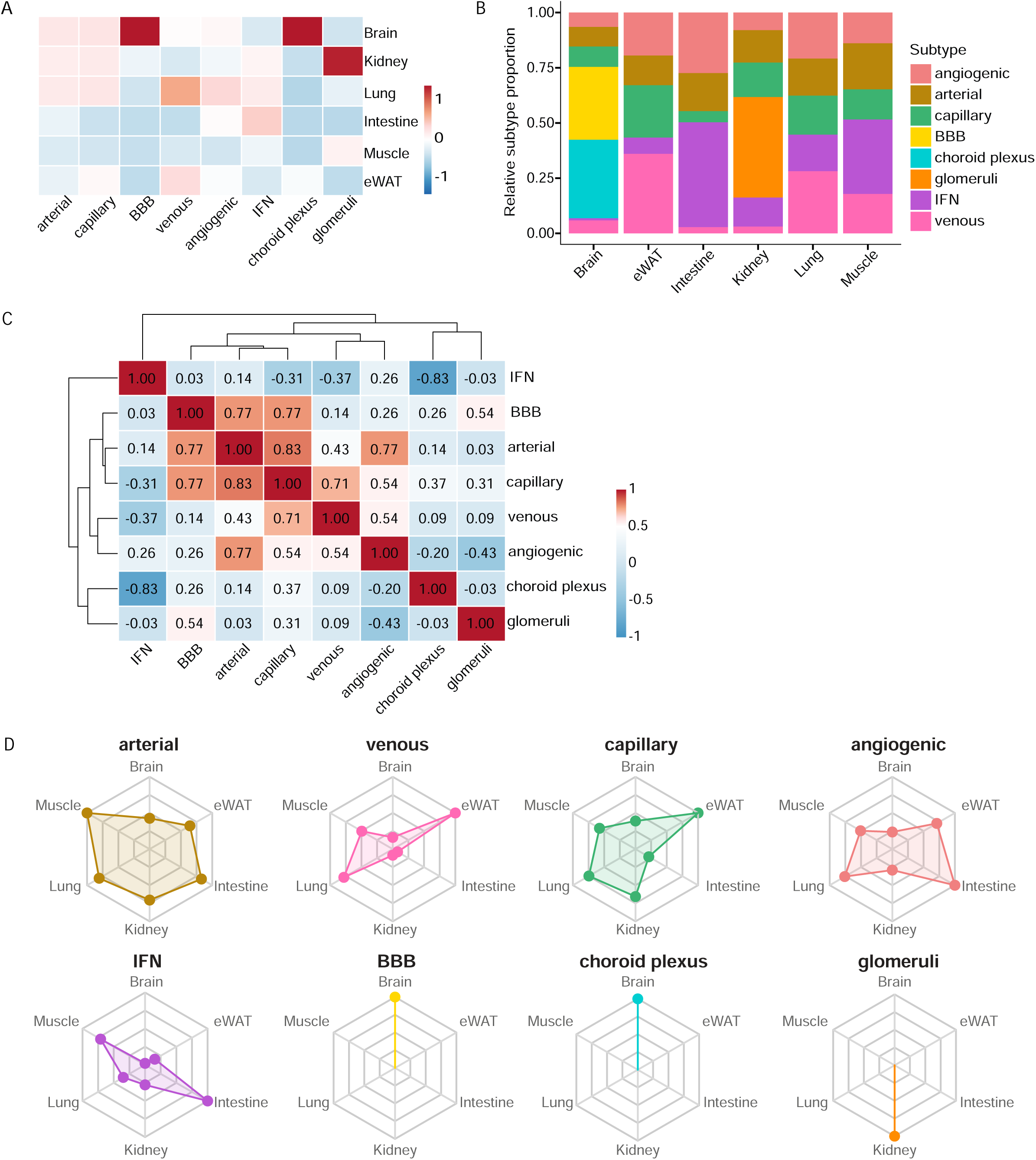
Proteomic deconvolution resolves endothelial subtype composition and tissue-specific zonation. **A,** Heatmap showing endothelial subtype scores across six tissues derived from bulk endothelial surface proteomes using z-score-based marker aggregation. **B,** Stacked bar plot depicting the relative composition of endothelial subtypes within each tissue based on z-score-based aggregation. **C**, Spearman correlation matrix of endothelial subtype scores across tissues, revealing structured relationships among vascular subtypes. **D,** Radar plots illustrating the distribution of individual endothelial subtypes across tissues.

Cross-method comparison showed that the z-score-based aggregation approach produced results that were broadly consistent with the other two deconvolution methods across endothelial subtypes (Supplementary Figure 4C). Agreement was strongest for BBB-like and choroid plexus endothelial signatures, whereas arterial, capillary, and angiogenic subtypes showed weaker and more variable correlations across methods. These findings indicate that the z-score-based approach provides a stable summary of subtype enrichment while preserving biologically meaningful differences across tissues. Using z-score-based marker aggregation, we next characterized endothelial subtype distributions across tissues (Figure 3B). Brain showed prominent BBB and choroid plexus signatures, whereas kidney showed a distinct glomerular endothelial component. In contrast, peripheral tissues displayed more distributed subtype profiles composed of canonical arterial, capillary, venous, angiogenic, and IFN-responsive signatures, without a single dominant specialized subtype. Notably, the intestine showed a relatively larger IFN-responsive component than most other tissues, potentially reflecting the immune-active mucosal environment of the gut. These distributions are consistent with organ-specific vascular specialization and indicate that *in vivo* surface proteomics captures broad endothelial subtype features across tissues. Spearman correlation analysis of subtype scores revealed coordinated relationships among endothelial subtype signatures (Figure 3C). Arterial, capillary, venous, and angiogenic signatures showed positive associations, consistent with partially shared marker structure among canonical endothelial programs. IFN-responsive, choroid plexus, and glomerular signatures showed more distinct correlation patterns, reflecting specialized or tissue-restricted endothelial states. Together, these analyses demonstrate that marker-based proteomic scoring captures both shared vascular programs and organ-specialized endothelial features. Finally, radar plot visualization provided an integrated view of endothelial subtype distributions across tissues (Figure 3D). Glomerular, BBB, and choroid plexus endothelial cells exhibited very distinct tissue-specificity, whereas arterial, capillary, and venous endothelial cells showed broad tissue distribution with organ-dependent gradients. Together, these findings demonstrate that proteomic deconvolution captures both classical and specialized endothelial states across vascular beds.

### Systematic, class-specific detection differences between CHRP surface proteomics and endothelial transcriptomics from scRNA-seq across organs

To systematically compare endothelial membrane proteins detected by *in vivo* CHRP-based surface proteomics and genes encoding endothelial membrane proteins detected by single-cell RNA sequencing (scRNA-seq) across tissues, we developed a cross-tissue enrichment framework integrating functional membrane protein classification, modality-specific gene set definition, and tissue-aware statistical analysis (Figure 4A). CHRP-derived endothelial surface proteomes from six tissues were integrated with endothelial transcriptomic references derived from two complementary single-cell RNA-seq atlases. For brain, kidney, lung, small intestine, and skeletal muscle, endothelial reference signatures were obtained from the Murine Endothelial Cell Atlas (E-MTAB-8077; Kalucka et al.). Endothelial populations in eWAT were independently defined using a dedicated single-cell atlas of human and mouse white adipose tissue (GSM5359340, GSM5359345, and GSM5359346). In this dataset, endothelial clusters were identified based on canonical endothelial markers, including *Pecam1*, *Cdh5*, *Kdr*, *Esam*, *Emcn*, and *Robo4* (Supplementary Figure 4A-B).

**Figure 4.**
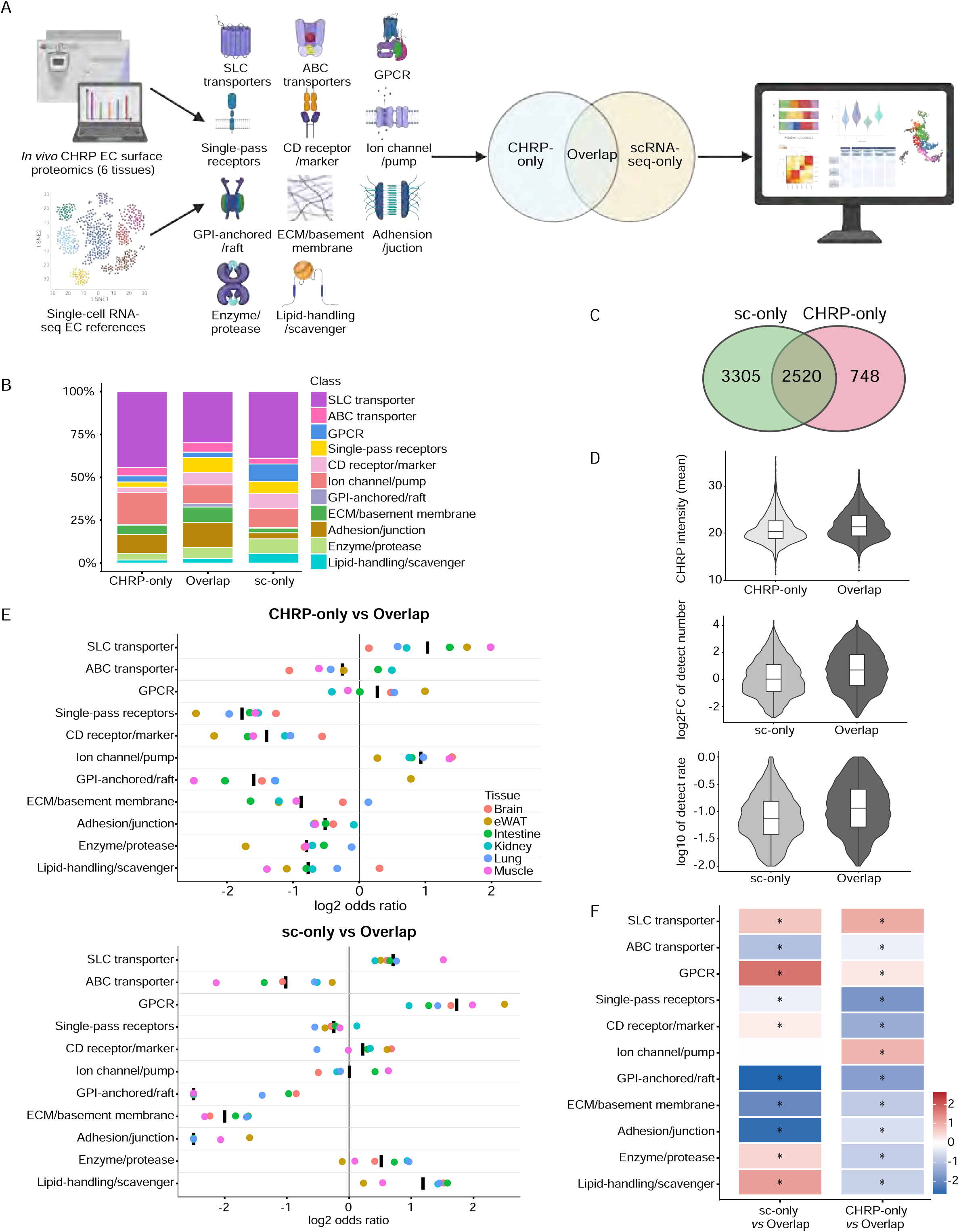
Cross-tissue enrichment analysis reveals modality-specific membrane protein class biases between CHRP proteomics and endothelial scRNA-seq. **A,** Schematic overview of the analytical framework used to compare endothelial membrane protein class representation between *in vivo* CHRP-based surface proteomics and endothelial single-cell RNA sequencing (scRNA-seq) references across six tissues. Membrane proteins were assigned to 11 functional classes based on curated Gene Ontology Cellular Component annotations. Within each tissue, proteins were independently categorized into CHRP-only, Overlap, or scRNA-seq-only (sc-only) sets based on detection by each modality. **B,** Stacked bar plots showing pooled (raw) membrane protein class composition for CHRP-only, Overlap, and sc-only sets aggregated across tissues. **C,** Venn diagram summarizing overlaps between endothelial membrane proteins detected by scRNA-seq (union across tissues) and CHRP proteomics (union across tissues). **D,** Violin plots showing pooled, cross-tissue comparisons of modality-specific detection metrics between CHRP-only vs overlap and scRNA-seq-only (sc-only) vs overlap protein sets. CHRP signal intensity is summarized as the log□-transformed mean CHRP intensity per protein. scRNA-seq detection is summarized using (i) detection breadth, defined as the log□ fold-change of the number of endothelial cells expressing each gene relative to the median value of scRNA-seq-only genes within each tissue (with a pseudocount of 1), and (ii) detection rate, defined as the fraction of endothelial cells expressing the gene and summarized as the log□□-transformed detection rate. All metrics were normalized within each tissue prior to pooling. Boxplots indicate median and interquartile range. **E,** Tissue-level enrichment of membrane protein classes across modalities. Dot plots show tissue-specific log□ odds ratios for enrichment of membrane protein classes in CHRP-only vs Overlap (top) and sc-only vs Overlap (bottom) comparisons across six tissues. Colored points represent individual tissue-level log□ odds ratios, while thick black bars indicate pooled, tissue-balanced log□ odds ratios across tissues. **F,** Heatmap summarizing pooled, tissue-balanced log_2_ odds ratios for membrane protein class enrichment in sc-only vs Overlap and CHRP-only vs Overlap comparisons. Stars indicate statistically significant enrichment after Benjamini-Hochberg correction (adjusted P < 0.05).

Endothelial membrane proteins identified in each tissue were assigned to 11 functional membrane protein classes using a curated, symmetric GO:Cellular Component-based annotation scheme. Within each tissue, proteins were independently categorized as CHRP-only, Overlap (detected by both modalities), or scRNA-seq-only (sc-only) (Figure 4A). The raw pooled membrane protein class composition across all tissues exhibited markedly distinct class distributions (Figure 4B). These differences reflected divergent representation of transporter-, receptor-, ECM-, and lipid-handling-associated classes between detection modalities, indicating that CHRP proteomics and scRNA-seq sample different regions of the endothelial membrane proteome. As raw pooled composition is inherently influenced by tissue-specific protein class prevalence, we next examined tissue-resolved membrane protein class composition across modalities (Supplementary Figure 4C, left panels). Across all six tissues, CHRP-only, Overlap, and sc-only sets showed reproducible yet tissue-dependent differences in class composition, indicating that modality-specific biases are present within individual tissues rather than arising solely from cross-tissue aggregation.

Comparison of endothelial membrane proteins detected by either modality revealed substantial but incomplete overlap. When aggregated across tissues, thousands of membrane protein genes were detected exclusively by scRNA-seq or exclusively by CHRP proteomics, with a large, shared set detected by both modalities (Figure 4C). Tissue-resolved Venn diagrams further demonstrated that the degree of overlap varied by tissue but consistently showed sizeable modality-specific gene sets (Supplementary Figure 4C, right panels). These results indicate that incomplete overlap between CHRP and scRNA-seq detection is a general feature across tissues rather than being driven by a single tissue context.

To quantify differences between modality-specific and overlapping proteins, we compared detection metrics across tissues. In pooled analyses, overlapping proteins consistently exhibited higher CHRP signal intensity compared with CHRP-only proteins. Similarly, scRNA-seq-derived metrics showed that sc-only proteins displayed reduced detection breadth and lower detection rates relative to the overlapping set (Figure 4D). Tissue-resolved violin plots demonstrated that these patterns were broadly consistent across all six tissues, despite variation in absolute signal magnitude and distribution shape (Supplementary Figure 4D). Effect sizes quantified using Cliff’s delta (Table S3) indicated that these differences represent systematic shifts in detectability rather than isolated outliers.

We next assessed tissue-specific enrichment of membrane protein classes using log□ odds ratios derived from Fisher’s exact tests (Figure 4E). Across tissues, multiple membrane protein classes showed consistent directional enrichment or depletion in CHRP-only or sc-only sets relative to overlapping proteins, although the magnitude of enrichment varied by tissue. Association strength, quantified using Cramér’s V (Table S4), indicated that these patterns reflect structured modality-dependent biases rather than random variation.

Because tissue-level analyses revealed consistent but heterogeneous class-level biases across organs (Figure 4E), we next asked whether these trends persisted after accounting for differences in tissue composition. To address this, we performed a pooled, tissue-balanced enrichment analysis that integrates information across all tissues while minimizing tissue-driven effects (Figure 5F). This analysis reveals robust and largely consistent functional class-level biases between CHRP-only and sc-only proteins relative to the overlapping set (Table S5). After tissue balancing, most membrane protein classes showed significant modality-specific enrichment or depletion after Benjamini-Hochberg correction. CHRP-only proteins were enriched for ion channels/pumps and SLC transporters but depleted for several receptor-associated, junctional, ECM/basement membrane, and raft-associated classes. Conversely, scRNA-seq-only proteins were enriched for GPCRs, lipid-handling/scavenger proteins, and enzymes/proteases, while showing depletion of structural and junctional classes. Ion channel-related proteins in the scRNA-seq-only set were the main class that did not reach significance after correction. Together, these analyses demonstrate that CHRP proteomics and endothelial scRNA-seq exhibit systematic, class-specific differences in membrane protein detection across tissues, extending beyond tissue-specific expression and reflecting modality-dependent biases in sampling the endothelial surface proteome.

**Figure 5.**
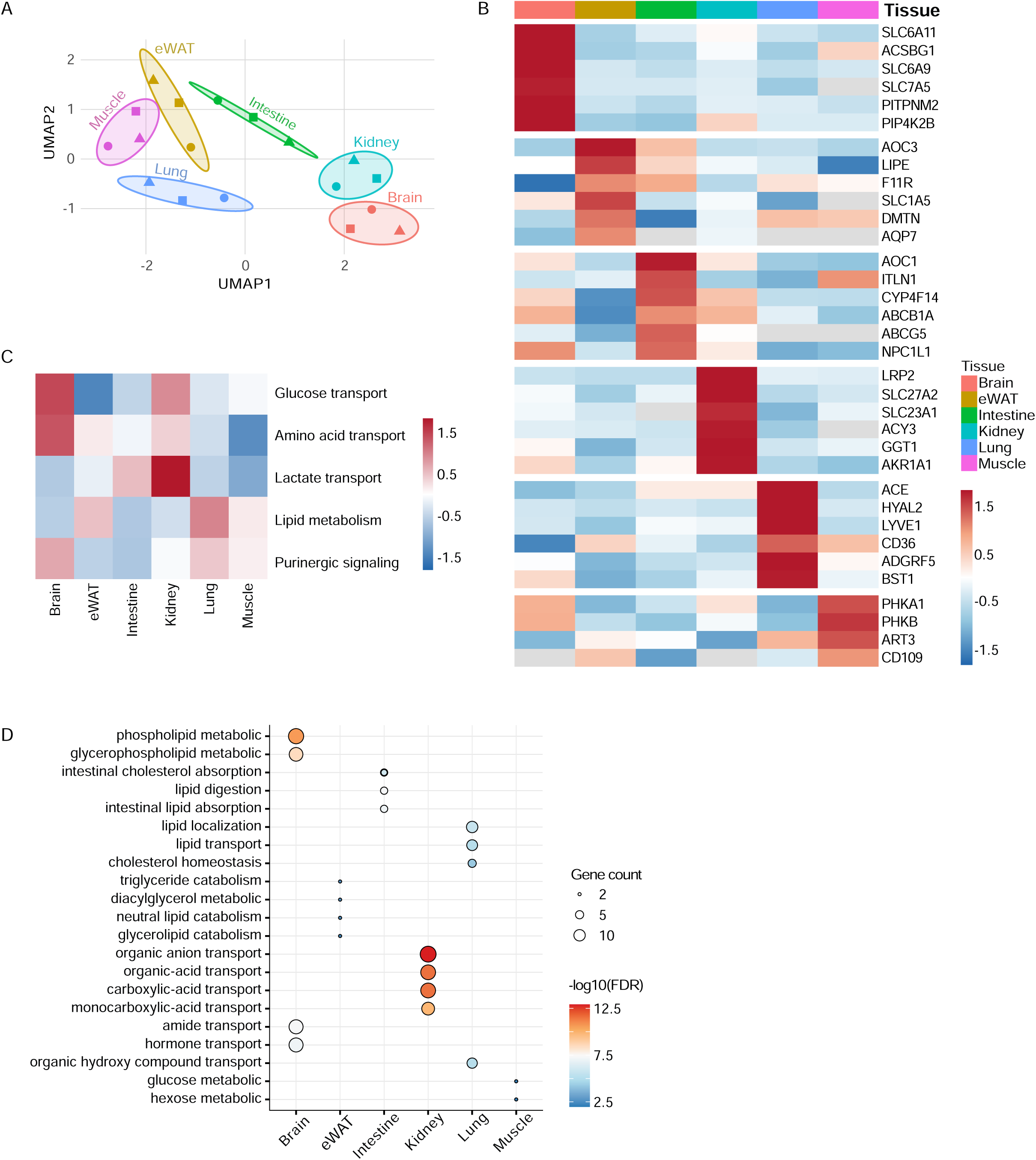
Endothelial surface proteomes define tissue-specific metabolic membrane proteins. **A,** UMAP visualization of EC surface metabolic membrane proteins across six tissues: brain, eWAT, intestine, kidney, lung, and skeletal muscle. Each point represents one biological replicate. **B,** Heatmap showing tissue-enriched EC surface metabolic membrane proteins across six tissues. Metabolic surface proteins were defined by membrane annotation and overlap with MSigDB metabolism-associated gene sets. Candidate proteins were ranked within each primary tissue using a composite score incorporating tissue enrichment, primary-tissue abundance, and replicate support. Up to the top six candidates per tissue are shown. Values represent row z-score-normalized protein abundance across tissues. **C,** Functional module heatmap summarizing major metabolic membrane programs across tissues. Metabolic membrane proteins were grouped into curated functional modules, including glucose transport, amino acid transport, lactate transport, lipid metabolism, and purinergic signaling. Colors represent scaled module scores across tissues. **D,** Gene Ontology enrichment analysis of tissue-assigned metabolic membrane proteins. Bubble plot showing selected metabolic and metabolite transport-related biological processes. Dot size indicates gene count, and color indicates -log_10_(FDR).

### Endothelial surface proteomes encode tissue-specific metabolic membrane signatures

Given the central role of endothelial cells in regulating nutrient exchange and metabolic homeostasis across vascular beds, we next asked whether endothelial surface proteomes contain tissue-specific metabolic membrane proteins involved in nutrient transport, metabolite exchange, and extracellular metabolic signaling. .

Dimensionality reduction analysis of these metabolic membrane proteins revealed clear segregation of biological replicates by tissue (Figure 5A), indicating that endothelial metabolic membrane proteomes differ across vascular beds. Consistent with this observation, hierarchical clustering of all membrane metabolic proteins showed marked tissue-dependent variation in protein abundance (Supplementary Figure 6A). Pairwise correlation analysis further confirmed divergent metabolic membrane profiles between tissues (Supplementary Figure 6B).

To identify tissue-enriched metabolic membrane proteins, we ranked candidates by tissue-selective abundance and focused on representative high-confidence proteins with consistent detection across biological replicates (Figure 5B). This analysis revealed distinct metabolic membrane programs across endothelial populations. Brain-enriched candidates were dominated by solute carriers and membrane-associated metabolic regulators, including SLC6A11, SLC6A9, SLC7A5, and PITPNM2. eWAT showed enrichment of lipid- and nutrient-handling proteins, including LIPE, AOC3, SLC1A5, AQP7, and PNPLA2. Intestinal candidates included proteins linked to lipid, cholesterol, xenobiotic, and nutrient handling, such as AOC1, CYP4F14, ABCB1A, ABCG5, and NPC1L1. Kidney-enriched proteins were strongly associated with solute, vitamin, fatty-acid, and glutathione-related metabolism, including SLC27A2, SLC23A1, GGT1, and AKR1A1. Lung and skeletal muscle candidates were more enriched for membrane-associated metabolic regulatory proteins, including vascular enzymes, lipid-handling proteins, and signaling-linked surface molecules. Together, these tissue-enriched candidates highlighted distinct endothelial metabolic interfaces across organs.

To further summarize functional organization, we grouped membrane metabolic proteins into curated modules representing glucose transport, amino acid transport, lactate transport, lipid metabolism, and purinergic signaling. Module-level scoring revealed tissue-biased metabolic membrane programs across vascular beds (Figure 5C). Brain showed prominent glucose and amino acid transport signatures, whereas kidney showed enrichment of lactate transport. Lung displayed stronger lipid metabolism and purinergic signaling-associated scores. Representative proteins underlying these module-level patterns are shown in Supplementary Figure 6C.

Finally, Gene Ontology enrichment analysis of tissue-assigned metabolic membrane proteins revealed distinct biological processes across vascular beds (Figure 5D). Brain candidates were enriched for phospholipid and glycerophospholipid metabolism and transport-related processes, whereas eWAT candidates were linked to lipid catabolism. Intestinal proteins were associated with cholesterol and lipid absorption, kidney proteins with organic anion and organic acid transport, and lung proteins with lipid localization, lipid transport, and cholesterol homeostasis. Muscle-associated candidates showed more limited enrichment, with terms related to glucose and hexose metabolism. Together, these analyses indicate that endothelial surface proteomes encode tissue-specific metabolic membrane programs that may support organ-specialized nutrient exchange, lipid handling, metabolite transport, and extracellular metabolic signaling.

### Identification of tissue-selective endothelial membrane protein candidates

Tissue-selective endothelial membrane proteins represent a candidate class for organ-specific vascular targeting. We therefore applied a strategy to define tissue-selective membrane candidates from this endothelial surface proteome atlas (Figure 6A). Candidate proteins were restricted to UniProt-annotated membrane-associated proteins, and 839 tissue-selective endothelial membrane candidates were identified across six tissue vascular beds (Table S6).

**Figure 6.**
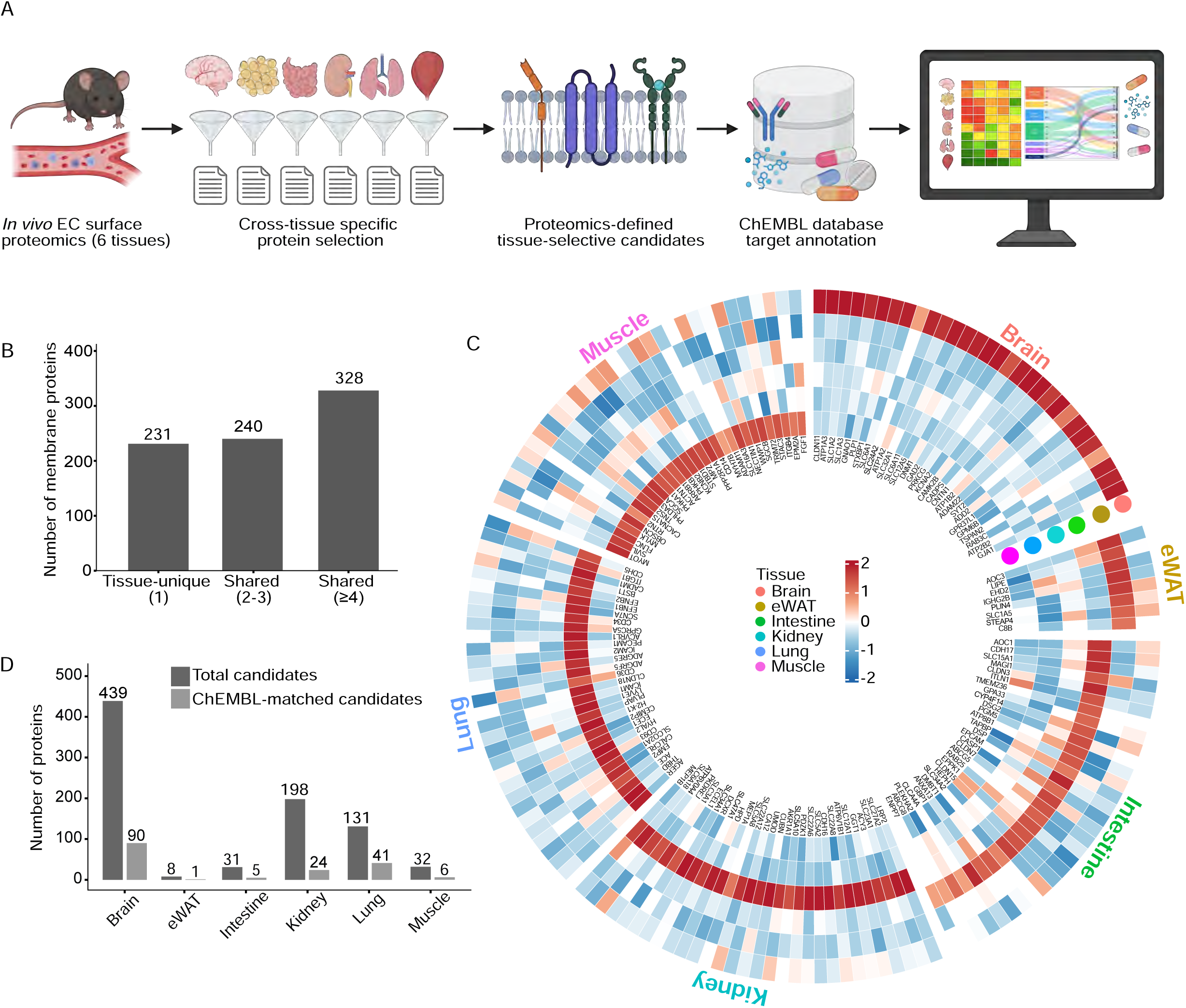
CHRP-defined tissue-selective endothelial membrane protein candidates inform organ-selective vascular target annotation. **A,** Workflow for proteomics-defined endothelial membrane candidate selection and target annotation. CHRP proteomic profiles from six tissues were used to identify tissue-selective endothelial membrane protein candidates. Candidate proteins were restricted to UniProt-annotated membrane proteins and selected based on tissue enrichment, primary-tissue abundance, and replicate detection support. Candidate proteins were then cross-referenced with the ChEMBL database to annotate existing compound-target relationships. **B,** Distribution of tissue-unique and shared CHRP-defined tissue-selective endothelial membrane protein candidates. Candidates were classified as tissue-unique, shared across 2-3 tissues, or shared across ≥4 tissues. **C,** Circular heatmap showing representative CHRP-defined tissue-selective endothelial membrane protein candidates across six tissues. For each tissue, up to top 30 prioritized candidates are shown. Candidates were prioritized using a composite proteomics-based score incorporating normalized tissue enrichment, normalized primary-tissue abundance, and replicate support. Values are shown as row-scaled z-scores across tissues. The complete tissue-selective endothelial membrane candidate list is provided in Table S6. **D,** ChEMBL annotation summary of CHRP-defined tissue-selective endothelial membrane protein candidates. Bars show the total number of membrane candidates identified in each tissue and the subset with ChEMBL compound-target annotations.

Quality-control analysis showed that the selected membrane candidates generally had strong signal in their primary tissue and were enriched relative to the second-highest tissue (Supplementary Figure 7A). We next assessed whether these candidates were tissue-restricted or shared across vascular beds, and identified 231 tissue-unique candidates, 240 candidates shared across 2-3 tissues, and 328 candidates shared across ≥4 tissues (Figure 6B). UpSet-like analysis further resolved the most frequent tissue-overlap patterns, showing both organ-restricted candidates and candidates shared by specific tissue groups (Supplementary Figure 7B). Together, these results suggest that vascular targeting candidates span a continuum from tissue-restricted to broadly shared membrane proteins.

To visualize representative candidates, proteins were ranked within each primary tissue using a composite proteomics-based score incorporating tissue enrichment, primary-tissue abundance, and replicate support. The top 30 prioritized candidates per tissue were displayed in a circular heatmap (Figure 6C). This analysis showed that the prioritized membrane candidates differed markedly across vascular beds, with brain, kidney, and lung contributing larger candidate groups, whereas eWAT, intestine, and skeletal muscle showed smaller and more selective candidate sets.

To further assess the translational potential of tissue-selective EC surface candidates, we cross-referenced the Table S6 candidate atlas with the ChEMBL 30 database, a manually curated drug discovery resource that integrates chemical, bioactivity, and genomic information to link molecular targets with bioactive compounds and therapeutic mechanisms. This analysis identified ChEMBL-matched candidates across multiple tissues, indicating that candidate proteins from the CHRP-derived tissue-selective surfaceome can be systematically queried against existing compound-target databases (Figure 6D). Representative tissue-target-compound relationships were further visualized by alluvial plot, highlighting candidate proteins and corresponding ChEMBL-annotated compounds across vascular beds (Supplementary Figure 7C). The full compound-target annotation output was compiled as Table S7.

To further characterize the functional classes represented in the membrane candidate atlas, we performed GO molecular function enrichment for all six tissues using tissue-selective membrane candidates as foreground and all CHRP-detected membrane proteins as background. Significant terms were detected in brain, kidney, and lung (Supplementary Figure 7D). These enriched functions aligned well with the expected biology of each vascular bed: brain candidates were associated with receptor, channel, and transporter activities; kidney candidates were dominated by active and secondary active transmembrane transporter functions; and lung candidates were linked to signaling receptor activity, cytokine and growth-factor binding, and adhesion-related functions. Together, these analyses establish an endothelial membrane candidate resource, linking ChEMBL-matched candidates to existing compound-target annotations while nominating the broader membrane candidate set for future organ-selective therapeutic target evaluation.

## DISCUSSION

Across vascular beds, endothelial cells line the luminal surface of blood vessels and form the primary interface between circulating blood, tissue microenvironments, and therapeutic agents. In this study, we established an endothelial-directed *in vivo* proximity-labeling strategy that combines genetically encoded membrane-tethered HRP with unbiased mass spectrometry to systematically map endothelial surface proteomes across tissues. This protein-level atlas resolves tissue- and subtype-associated endothelial surface features, reveals differences between proteomic and transcriptomic detection of endothelial membrane proteins, and nominates tissue-selective endothelial membrane candidates for organ-selective vascular targeting. Our work bridges a critical gap between transcriptional atlases of vascular heterogeneity and the endothelial surface protein landscape that shapes vascular accessibility, signaling, and future therapeutic targetability.

A major insight from this study is that endothelial transcriptomic and surface proteomic profiles capture complementary, but not interchangeable, aspects of vascular biology. The discordance observed here was not limited to individual targets, but reflected systematic, class-specific differences in endothelial membrane protein representation across tissues. By comparing CHRP-based surface proteomics with endothelial single-cell transcriptomic references, we found that distinct classes of membrane proteins were preferentially detected by each modality in a reproducible and tissue-consistent manner. These findings suggest that scRNA-seq and *in vivo* surface proteomics sample different layers of the endothelial surfaceome, shaped by transcriptional regulation, protein stability, membrane accessibility, recycling, and turnover [13, 40, 41]. Consequently, reliance on transcriptomic data alone may bias vascular target discovery toward RNA-detectable protein classes while underrepresenting stable, recycled, or post-transcriptionally regulated surface proteins [14, 42]. TFRC illustrates this principle. Although transferrin receptor has been widely explored in receptor-mediated BBB transport strategies [43–45], its transcript abundance was minimal in kidney endothelial populations in the Murine Endothelial Cell Atlas (E-MTAB-8077; Kalucka et al.). To exclude dataset-specific technical limitations, we also interrogated an independent kidney single-cell RNA-seq dataset (GSE107585) [46], which similarly revealed *Tfrc* expression in fewer than 1% of endothelial cells (data not shown). In contrast, TFRC protein was robustly detected by *in vivo* CHRP proteomics among kidney-associated endothelial membrane candidates. This discrepancy is consistent with the biology of TFRC as a stable, highly recycled membrane proteins governed by endocytic trafficking pathways [47–50], which can maintain abundant surface pools without sustained high transcriptional activity. Such examples underscore the value of protein-level measurements in native tissue context for identifying vascular surface proteins that may be underestimated by transcriptomic approaches alone [51].

These modality-specific differences also shaped how we interpreted endothelial heterogeneity from CHRP proteomic profiles. A key analytical consideration was how to extract endothelial subtype information from bulk *in vivo* proteomic measurements without overinterpreting the data as quantitative cell-type proportions. Unlike transcriptomic deconvolution, proteomic inference is constrained by incomplete protein coverage, non-amplifiable signals, missing values, protein turnover, recycling, and labeling efficiency. We therefore interpreted endothelial subtype signatures as relative marker-enrichment patterns rather than direct estimates of subtype abundance. Among the approaches evaluated, z-score-based marker aggregation produced the most anatomically coherent endothelial subtype patterns in our dataset and was used to summarize arterial, venous, capillary, and tissue-specialized endothelial programs.

Beyond vascular subtype heterogeneity, our analyses revealed evidence of tissue-dependent metabolic specialization at the endothelial surface. Distinct vascular beds preferentially enriched membrane proteins involved in nutrient transport, lipid handling, organic acid transport, or glucose metabolism, suggesting that endothelial cells are functionally adapted to the metabolic demands of their surrounding tissue environment, consistent with the concept of organotypic vascular specialization described in transcriptomic studies of the vasculature [10, 52]. These findings extend previous transcriptomic observations of endothelial metabolic heterogeneity by demonstrating that such specialization is also reflected at the surface proteome level, where proteins are positioned to directly regulate exchange between the circulation and individual organs and contribute to tissue metabolic homeostasis [53, 54].

The same principle also guided our strategy for tissue-selective candidate discovery. By focusing on membrane-associated proteins, the candidate atlas emphasizes proteins most likely to be accessible to circulating ligands, antibodies, or engineered delivery systems. Although ChEMBL annotation provides one route to connect these candidates with existing compound-target knowledge, the broader value of Table S6 extends beyond targets already represented in chemical databases. Many tissue-enriched vascular membrane proteins without current ChEMBL matches may still provide useful starting points for antibody development, ligand discovery, disease-context validation, or future tissue-selective delivery strategies. SLC7A5 illustrates this point. SLC7A5 was retained in Table S6 as a brain-enriched endothelial membrane candidate and showed greater tissue selectivity than SLC3A2 (CD98hc), its binding partner and a BBB transport target that has been explored for brain delivery [55]. SLC7A5 (LAT1) is a major amino acid transporter at the blood-brain barrier [56, 57] and forms a functional heterodimer with SLC3A2 [58]. This example suggests that the atlas may help nominate tissue-enriched components of transporter complexes for future BBB-targeting strategies, rather than focusing only on targets with existing compound annotations or single surface markers.

Recent work using lectin-conjugated peroxidase perfusion has similarly underscored the value of *in vivo* luminal surface proteomics for BBB biology, revealing the mouse brain vascular luminal surface proteome and identifying BBB regulators such as SLC7A1 and HYAL2 [59]. This perfusion-based strategy is particularly well suited for barrier-forming vascular beds such as the brain, where tight junctions restrict probe access primarily to the luminal endothelial surface. In more permeable tissues, however, perfused probes may access extravascular surfaces and capture a broader perfusion-accessible proteome. CHRP provides a complementary strategy by genetically restricting HRP catalytic activity to defined cell types, thereby allowing probe access and HRP localization together to determine the labeled surface compartment. This principle was supported by our adipocyte-directed HRP experiment in adipose tissue, where vascular permeability permits labeling substrates to reach parenchymal cell surfaces. Adipose tissue is characterized by high vascular permeability, extensive plasma protein exchange, and close coupling between ECs and neighboring adipocytes [60–62]. In this setting, adipocyte-directed HRP produced selective adipocyte surface labeling, supporting the feasibility of genetically directed *in vivo* proximity labeling beyond ECs in tissues with permissive vascular access. Thus, while the central focus of this study is tissue-resolved endothelial surface proteomics, the FHRP experiment provides a proof of principle that the same labeling chemistry can be redirected to additional cell types when substrate delivery is supported by local vascular permeability.

Together, our findings establish HRP-mediated *in vivo* proximity labeling as a versatile approach for surface proteomics that can be tailored to distinct vascular barrier properties and biological questions. By capturing surface-exposed proteins *in vivo* within native tissue context, this approach provides a protein-level framework for defining vascular heterogeneity, evaluating target accessibility, and informing organ-selective therapeutic strategies.

## Figure legend

**Supplementary Figure 1.**
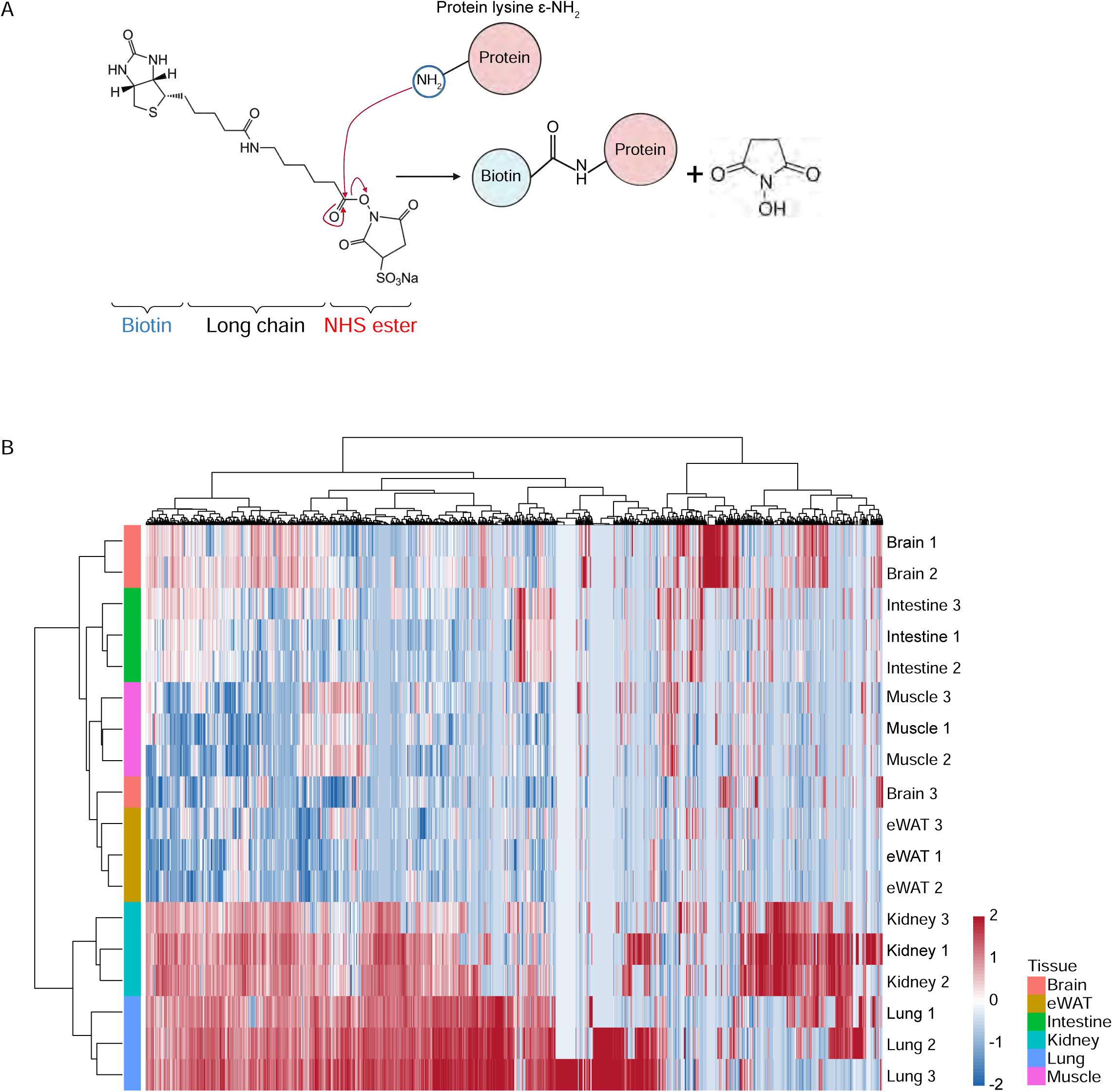
A, Chemical principle of Sulfo-NHS-Biotin labeling. Sulfo-NHS-Biotin reacts with accessible protein lysine ε-amine groups via nucleophilic attack on the activated NHS ester carbonyl of the biotin moiety, resulting in the formation of a stable amide linkage and covalent biotin conjugation. **Supplementary Figure 1B,** Sample-level heatmap showing NHS-Biotin-labeled proteins across individual biological replicates. Protein intensities were log□-transformed and Z-score normalized across samples for each protein. Hierarchical clustering was performed on individual samples using correlation distance and on proteins using Euclidean distance, with Ward’s linkage. Z-scores were capped at ±2 for visualization.

**Supplementary Figure 2.**
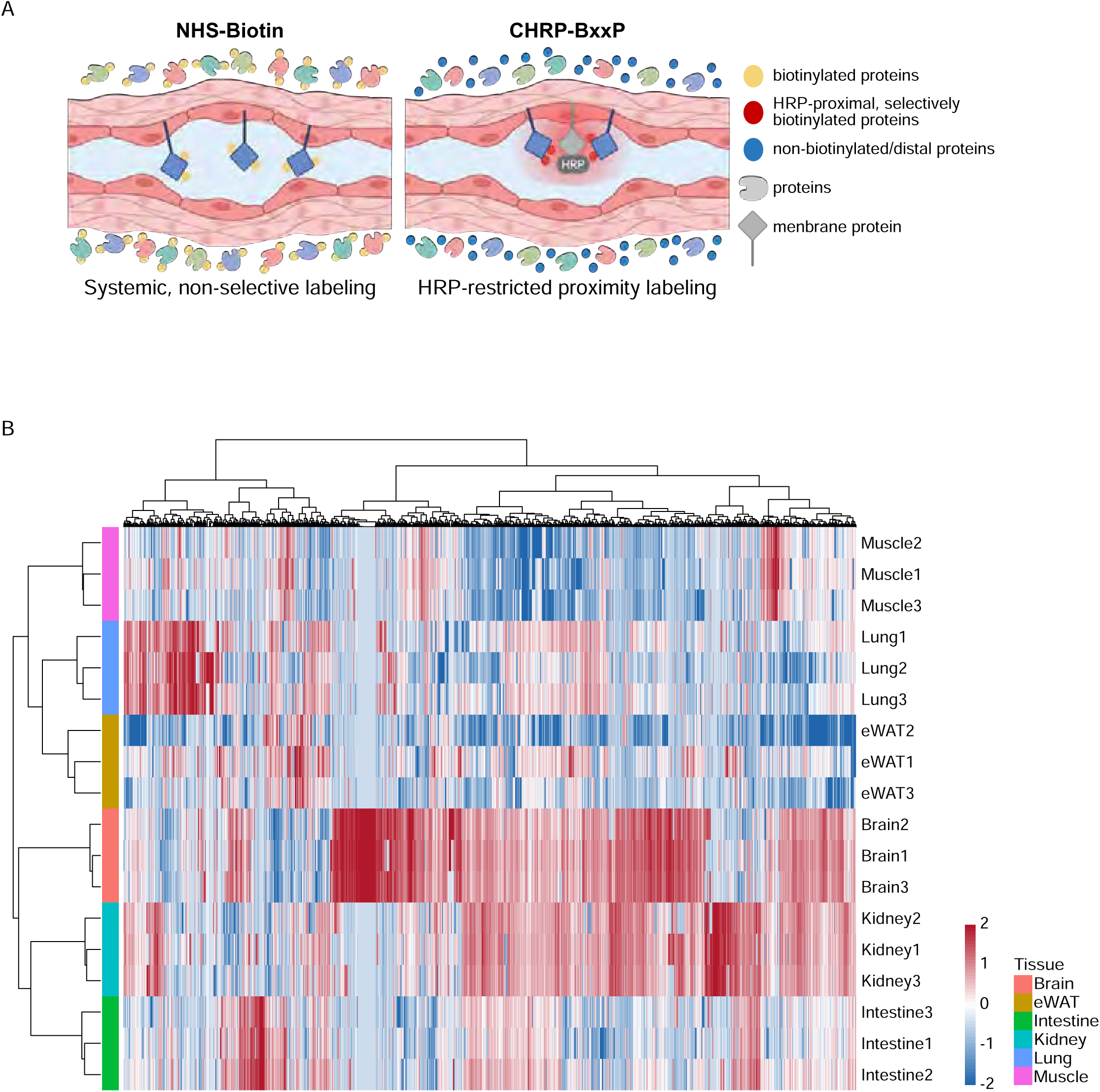
A, Schematic illustration comparing the principles of systemic NHS-Biotin labeling and CHRP-BxxP-based proximity labeling for *in vivo* biotinylation of vascular membrane proteins. Left, systemic perfusion of NHS-Biotin results in non-selective, amine-reactive labeling of accessible proteins throughout the vascular lumen, independent of spatial proximity or enzymatic localization. Right, in CHRP mice, HRP is specifically expressed at the endothelial luminal surface and catalyzes the oxidation of BxxP in the presence of hydrogen peroxide, generating short-lived reactive intermediates that covalently label proteins in close proximity to HRP. Proteins within the HRP-restricted labeling radius are selectively biotinylated, whereas distal proteins remain unlabeled, conferring spatially confined and tissue-specific labeling. **Supplementary Figure 2B,** Sample-level heatmap of CHRP-labeled proteins across individual biological replicates, with log□-transformed intensities and Z-score normalized across samples. Hierarchical clustering was performed on individual samples using correlation distance and on proteins using Euclidean distance, with Ward’s linkage. Z-scores were capped at ±2 for visualization.

**Supplementary Figure 3.**
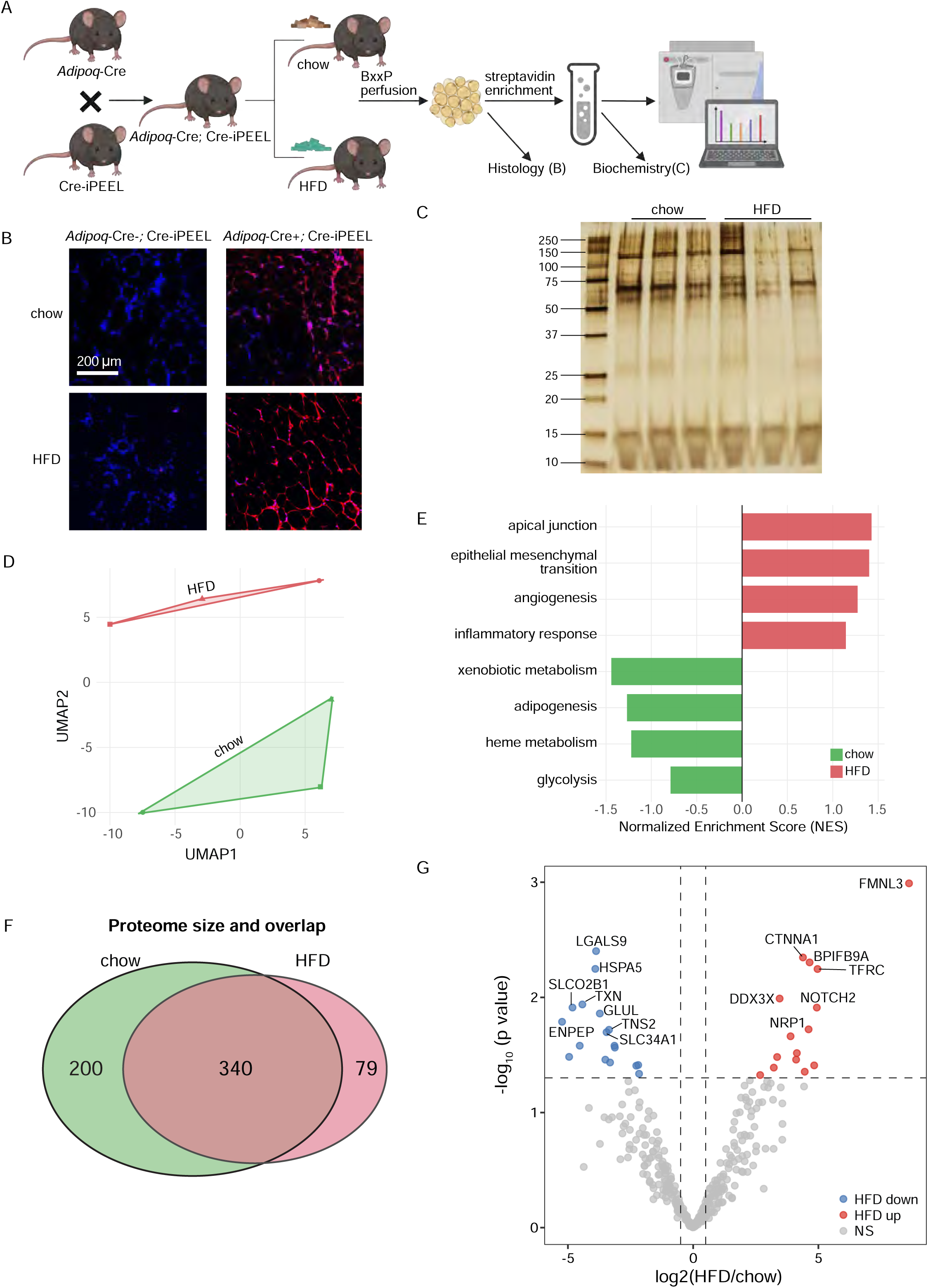
Adipocyte-restricted HRP-mediated *in vivo* proximity labeling captures diet-associated surfaceome changes. **A**, Mating scheme used to generate adipocyte-specific membrane-HRP mice, referred to as FHRP mice, for proof-of-principle *in vivo* proximity-labeling experiments. **B**, Representative immunofluorescence images showing adipocyte-surface biotinylation in chow and HFD FHRP mice following BxxP perfusion, visualized by Alexa Fluor 647-conjugated streptavidin. Scale bar=200 μm. **C,** Silver staining of streptavidin-enriched fractions from chow and HFD eWAT biological replicates. **D**, UMAP visualization of FHRP adipocyte surfaceome proteomic profiles from chow and HFD mice. **E**, Hallmark gene set enrichment analysis based on proteins ranked by HFD-versus-chow differential abundance, using MSigDB Hallmark gene sets obtained through msigdbr. Pathways enriched in HFD or chow samples are shown. **F**, Venn diagram showing shared and diet-associated adipocyte surface proteins identified in chow and HFD groups. **G**, Volcano plot of differential adipocyte surface proteins between HFD and chow groups, showing log□ fold change (HFD/chow) and -log□□ P value. Selected representative proteins are annotated.

**Supplementary Figure 4.**
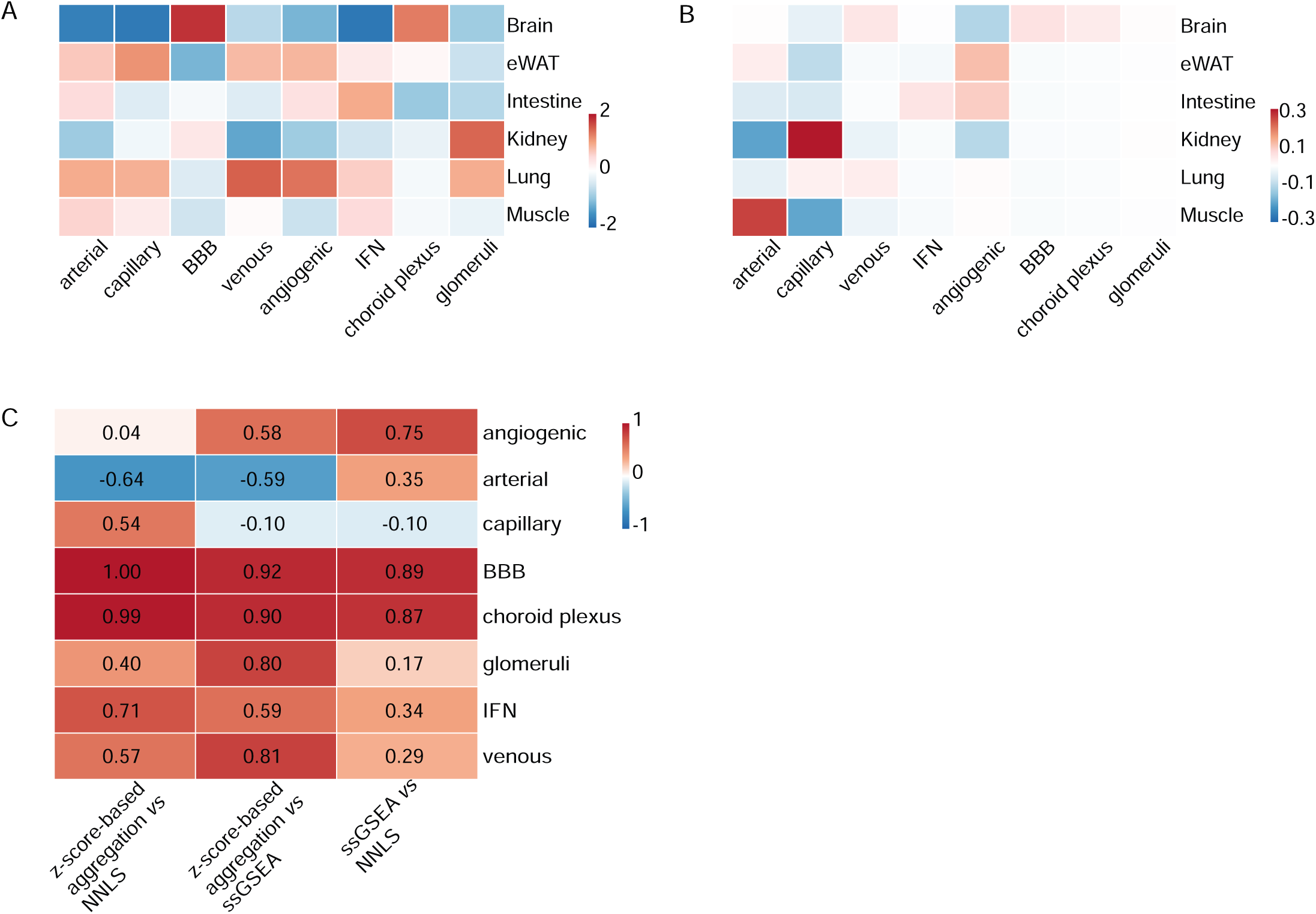
Comparison of proteomic deconvolution strategies. **A,** Endothelial subtype score heatmap generated using ssGSEA. **B,** Endothelial subtype score heatmap generated using non-negative least squares (NNLS). **C,** Pearson correlation analysis comparing subtype scores derived from z-score-based aggregation, ssGSEA, and NNLS.

**Supplementary Figure 5.**
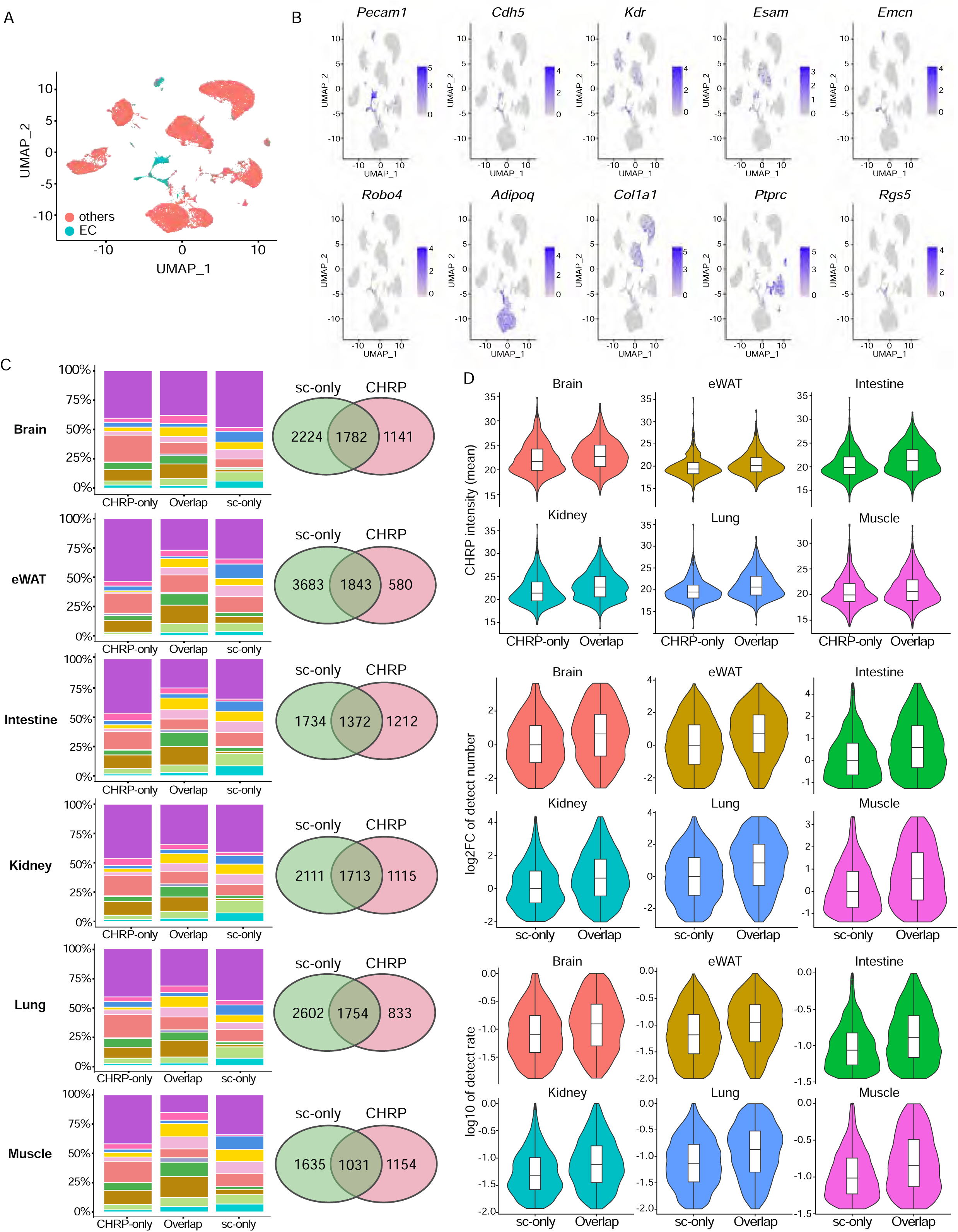
Tissue-resolved membrane protein class composition and modality-specific detection metrics. **A,** Single-cell clustering of eWAT identifying endothelial populations. **B,** Feature plots of canonical endothelial markers confirming EC identity. **C,** Tissue-specific membrane protein class composition and modality overlap. Left panels show stacked bar plots of the relative composition of 11 membrane protein classes for CHRP-only, Overlap, and sc-only gene sets within each tissue, based on a curated Gene Ontology Cellular Component-based classification scheme. Right panels show tissue-specific Venn diagrams summarizing overlap between endothelial membrane proteins detected by scRNA-seq and CHRP proteomics within each tissue. Numbers indicate unique and shared genes per modality. **D,** Tissue-resolved comparisons of modality-specific detection metrics corresponding to the pooled analyses in Figure 4D. Top row: CHRP signal intensity summarized as log□-transformed mean CHRP intensity comparing CHRP-only vs overlap proteins within each tissue. Middle row: scRNA-seq detection breadth defined as the log□ fold-change of the number of endothelial cells expressing each gene relative to the median value of scRNA-seq-only genes within each tissue (with a pseudocount of 1). Bottom row: scRNA-seq detection rate summarized as the log_10_-transformed fraction of endothelial cells expressing the gene. Boxplots indicate median and interquartile range.

**Supplementary Figure 6.**
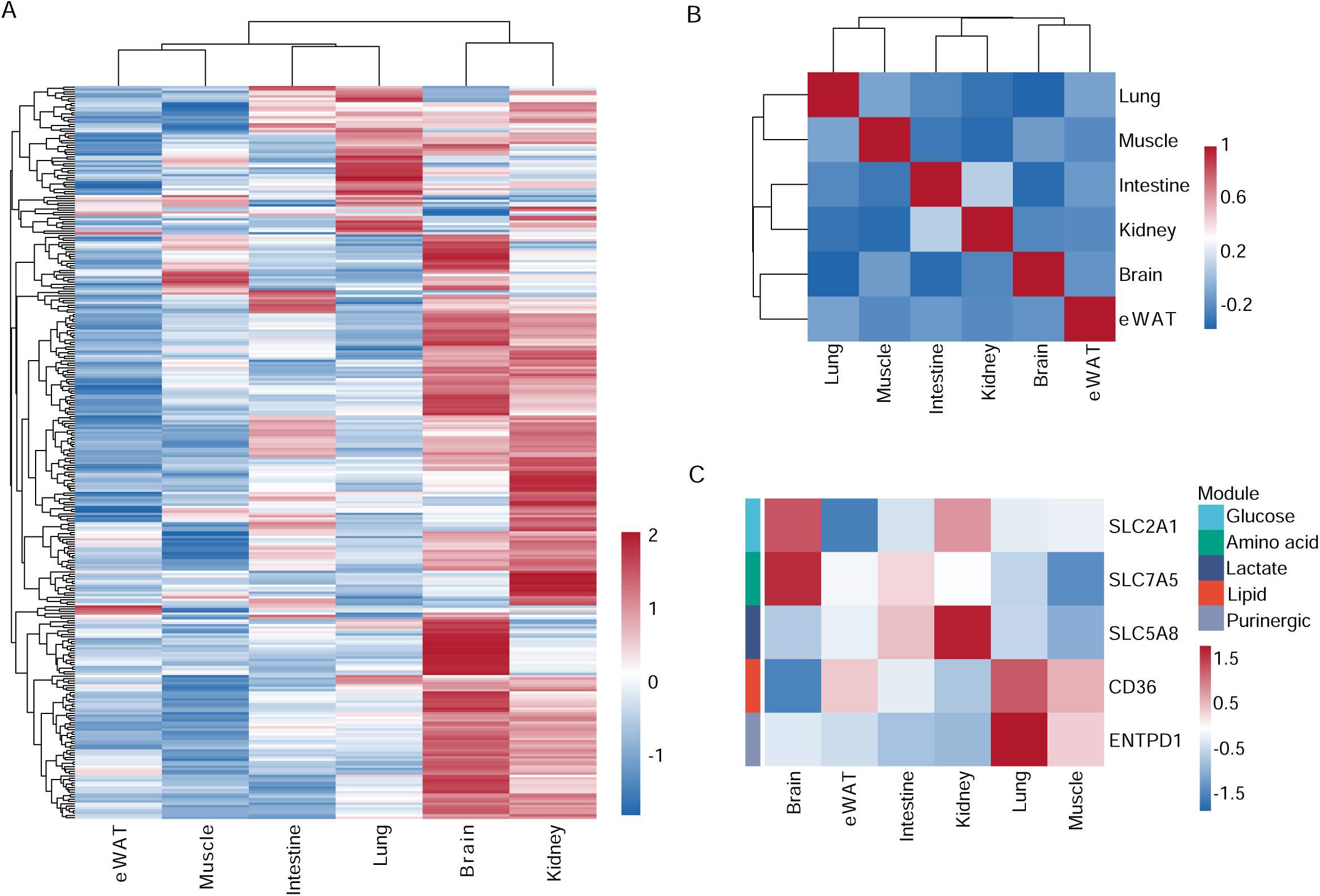
Supporting analyses of tissue-specific endothelial metabolic surface proteomes. **A**, Hierarchical clustering heatmap of all metabolic membrane proteins across tissues. Values represent row z-score-normalized tissue-averaged protein abundance. **B,** Pearson correlation heatmap of metabolic surface membrane proteomes between tissues based on tissue-level mean protein abundance profiles. **C,** Heatmap of representative metabolic surface membrane proteins underlying the functional modules shown in Figure 5C, illustrating selected drivers of tissue-specific metabolic membrane programs.

**Supplementary Figure 7.**
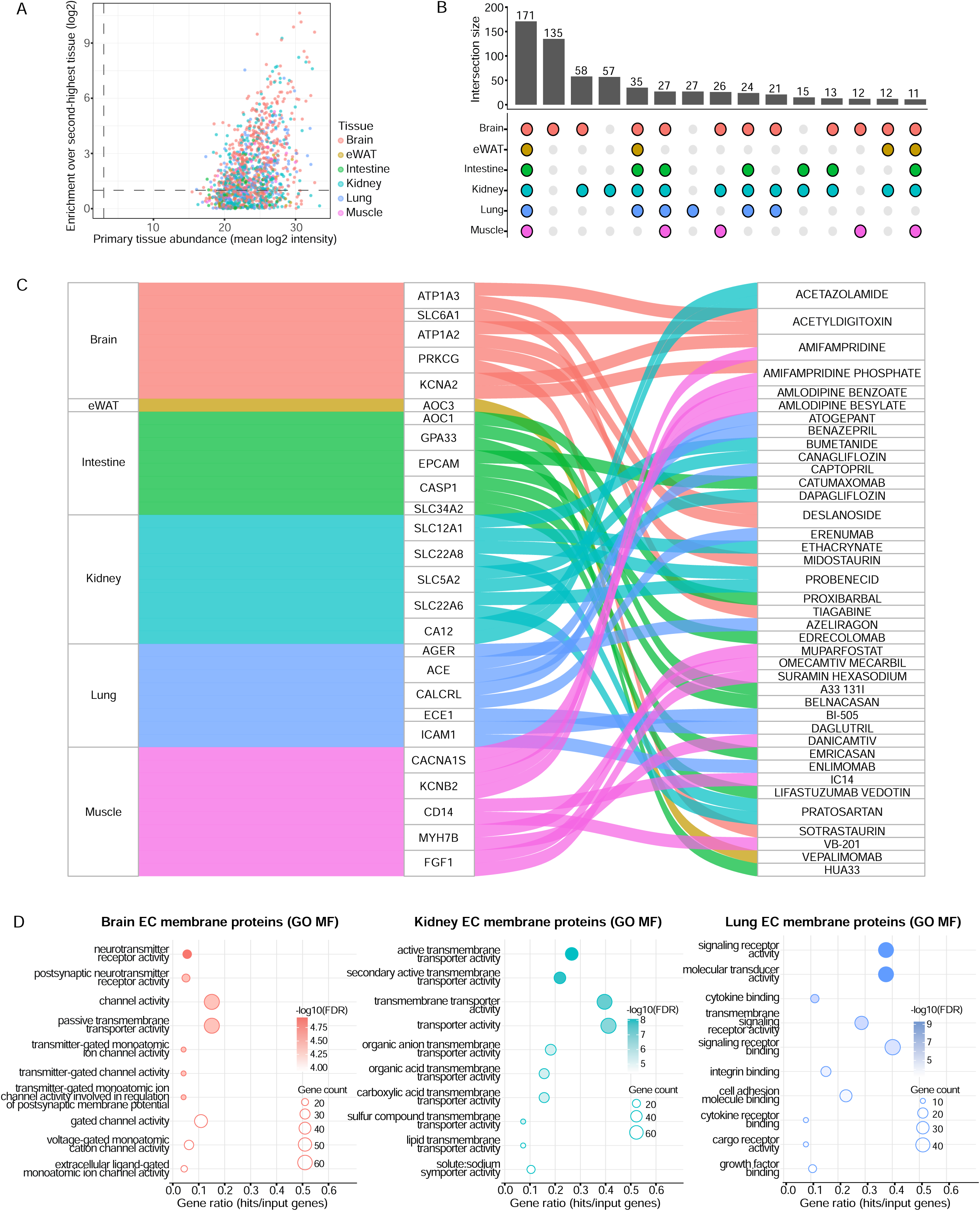
Supporting analyses for CHRP-defined tissue-selective endothelial membrane candidate selection and ChEMBL annotation. **A,** Candidate selection QC for CHRP-detected membrane proteins. Each dot represents one CHRP-detected membrane protein. The x-axis shows mean log_2_ abundance in the primary tissue, and the y-axis shows enrichment over the second-highest tissue. Dashed lines indicate the abundance and enrichment thresholds used for tissue-selective candidate selection. **B,** UpSet-like analysis showing exact tissue intersections among CHRP-defined tissue-selective endothelial membrane protein candidates. **C,** Alluvial plot showing representative ChEMBL compound-target annotations for CHRP-defined tissue-selective endothelial membrane candidates. Flows connect tissue of enrichment, candidate target proteins, and annotated ChEMBL compounds. For visualization, up to five ChEMBL-matched candidate proteins per tissue and up to two compounds per candidate protein are shown; the full compound-target annotation is provided in Table S7. **D,** GO molecular function enrichment analysis of CHRP-defined tissue-selective endothelial membrane protein candidates. Enrichment analysis was performed for all six tissues using tissue-selective membrane candidates as foreground and all CHRP-detected membrane proteins as background. Dotplots are shown for tissues with significantly enriched terms under the applied cutoffs, including brain, kidney, and lung. Dot size indicates gene count, and color indicates -log_10_(FDR).

