## Supplementary Methods for "A tissue-resolved endothelial surface proteome atlas informs organ-selective vascular targeting"

**Mice**

Animal care and experimental protocols were approved by the Institutional Animal Care and Use Committee of the Baylor College of Medicine. Mice were maintained under specific pathogen-free conditions on a 12-hour light/dark cycle with ad libitum access to food and water unless otherwise indicated.

To achieve endothelial cell–restricted expression of membrane-tethered horseradish peroxidase (HRP), *Cdh5*-CreERT mice, originally described by Wang et al.[1], were crossed with Cre-iPEEL mice (JAX #037698) to generate *Cdh5*-CreERT; Cre-iPEEL offspring (CHRP mice). For tamoxifen-induced recombination, tamoxifen (Sigma-Aldrich, T5648-1G) was dissolved in corn oil and administered by intraperitoneal injection (75 mg/kg/day) for 5 consecutive days. Mice were allowed a washout period of at least 7 days following the final injection prior to *in vivo* labeling experiments.

To assess non-selective vascular surface labeling, wild-type C57BL/6J mice (JAX #000664) were used for Sulfo-NHS-LC-Biotin (NHS-Biotin; ApexBio, A8003) perfusion experiments.

For adipocyte-specific surface proteomics, *Adipoq*-Cre mice (JAX #028020) were crossed with Cre-iPEEL mice to generate *Adipoq*-Cre; Cre-iPEEL offspring (FHRP mice). These mice were fed either a standard chow diet or a 60% high-fat diet (HFD; BioServ, S1850) for 15 weeks prior to *in vivo* proximity labeling and tissue collection.

**NHS-Biotin transcardial perfusion**

For non-selective *in vivo* vascular labeling, mice were anesthetized and perfused with NHS-Biotin dissolved in phosphate-buffered saline (PBS) at 0.5 mM for 10 minutes and then quenched by perfusion 50 mM glycine (Sigma-Aldrich, G7126-500G) in PBS for 3 minutes, and tissues were immediately harvested or further processed as indicated.

**HRP-mediated *in vivo* proximity labeling**

For cell-type-restricted proximity labeling, mice were perfused with Biotin-XX Tyramide (BxxP; (ApexBio Technology, A8012) as a labeling substrate. After an initial Tyrode’s buffer (145 mM NaCl, 1.25 mM CaCl_2_, 3 mM KCl, 1.25 mM MgCl_2_, 0.5 mM NaH_2_PO_4_, 10 mM glucose, 10 mM HEPES, pH 7.4) perfusion to clear blood, mice were perfused with BxxP (100 µM) dissolved in Tyrode’s buffer for 5 minutes to allow substrate distribution. Labeling was initiated by perfusion with BxxP plus hydrogen peroxide (0.003% H₂O₂) for 5 minutes,

and then terminated by immediate perfusion with quenching buffer (Tyrode’s buffer containing 10 mM sodium azide, 10 mM sodium ascorbate, and 5 mM Trolox) for 3 minutes. Following quenching, tissues were excised, snap-frozen in liquid nitrogen, or fixed for downstream analyses.

**Tissue lysis**

Biotinylated proteins were enriched from labeled tissues using streptavidin-based affinity purification. Briefly, 400 µl 1% SDS RIPA buffer (50 mM Tris-HCl (pH 8.0), 150 mM NaCl, 1% sodium dodecyl sulfate (SDS), 0.5% sodium deoxycholate (Thermo Scientific, 89904), 1% Triton X-100, 1x Protease and Phosphatase Inhibitor Cocktail (Thermo Scientific; 78442), and 1 mM phenylmethylsulfonyl fluoride (PMSF) (Thermo Scientific, 36978)) was added to 200 mg epididymal white adipose tissue (eWAT), or 40 mg other tissues (lung, kidney, skeletal muscle, small intestine and brain), and homogenized with TissueLyser (Qiagen, 9003240) for 3 minutes. Samples were then vortexed briefly, followed by two rounds of sonication at 4 ℃ (30% pulse, 15s, Qsonica sonicators) until the lysate became clear. Then samples were heated to 95℃ for 5 minutes and return onto ice for 1 minute. 1600 µl SDS-free RIPA buffer was added to each sample to yield 0.2% SDS normal RIPA buffer. Samples were cleared by centrifugation at 16,000 g for 15 minutes at 4℃.

**Streptavidin enrichment and mass spectrometry**

Streptavidin magnetic beads (Thermo Scientific, 88817) were used to enrich biotinylated proteins. After protein concentration was determined by Pierce™ BCA Protein Assay Kits (Thermo Scientific, 23227), the estimated amount of biotinylated protein for enrichment was calculated according to the manufacturer’s guidelines (each 100μL of streptavidin magnetic beads can capture approximately 55μg of biotinylated rabbit IgG). Protein lysates were subsequently incubated with streptavidin magnetic beads at 4 °C for 2 hours with gentle rotation. The beads were then sequentially washed twice with 1 mL normal RIPA buffer, once with 1 mL 1 M KCl, once with 1 mL 0.1 M Na_2_CO_3_, once with 1 mL 2 M urea in 10 mM Tris-HCl (pH 8.0), and twice with 1 mL normal RIPA buffer.

For silver staining, biotinylated proteins were eluted by heating the beads at 95 ℃ for 10 min in 50 µl 3x protein loading buffer supplemented with 10% 2-Mercaptoethanol and 2 mM biotin and then separated by SDS-PAGE and visualized using the Pierce™ Silver Stain for Mass Spectrometry kit (Thermo Scientific, 24600) according to the manufacturer’s instructions.

For proteomic analysis, streptavidin-enriched proteins underwent on-bead digestion overnight with 1 mg of trypsin (Thermo Scientific, 90059) following reduction with dithiothreitol (DTT) and alkylation with iodoacetamide (Sigma-Aldrich, I1149-5G). The resulting peptides were purified by solid-phase extraction using an Oasis HLB plate (Waters). LC-MS/MS analysis was performed at the Proteomics Core Facility of the University of Texas Southwestern Medical Center using a Q Exactive HF mass spectrometer coupled to an Ultimate 3000 RSLC-Nano liquid chromatography system. Peptides were loaded onto a 75-μm internal diameter × 15-cm EasySpray column (Thermo Scientific, ES900) and eluted using a 90-min gradient from 0% to 28% buffer B. Buffer A consisted of 2% (v/v) acetonitrile and 0.1% formic acid in water, whereas buffer B consisted of 80% (v/v) acetonitrile, 10% (v/v) trifluoroethanol, and 0.1% formic acid in water. The mass spectrometer was operated in positive-ion mode with a source voltage of 2.5 kV and an ion transfer tube temperature of 300 °C. MS scans were acquired at 120,000 resolution in the Orbitrap and up to 20 MS/MS spectra were obtained in the ion trap for each full spectrum acquired using higher-energy collisional dissociation (HCD) for ions with charges 2-8. Dynamic exclusion was set for 20 s after an ion was selected for fragmentation.

Raw mass spectrometry data were analyzed using Proteome Discoverer v3.0 (Thermo Scientific). Peptide-spectrum matching was performed using Sequest HT against the reviewed *Mus musculus* UniProt protein database. Precursor and fragment mass tolerances were set to 10 ppm and 0.02 Da, respectively, and up to three missed cleavages were permitted. Carbamidomethylation of cysteine was specified as a fixed modification, and oxidation of methionine was specified as a variable modification. INFERYS and Percolator were used within Proteome Discoverer for post-search rescoring using default settings. Peptide identifications were filtered at a false discovery rate (FDR) of 1%. Protein abundance was calculated as the summed peak intensities of all peptides assigned to each protein.

**Immunofluorescence**

To visualize *in vivo* proximity labeling and assess cellular and anatomical specificity, immunofluorescence staining was performed on tissue sections following perfusion-based labeling. After completion of NHS-Biotin or HRP-mediated proximity labeling, tissues were immediately excised and fixed overnight in 10% phosphate-buffered formalin. Fixed tissues were dehydrated, embedded in paraffin, and sectioned at the Pathology Core Facility at Baylor College of Medicine.

Paraffin sections were subjected to standard deparaffinization and rehydration procedures, followed by antigen retrieval in 10 mM sodium citrate buffer (pH 6.0) (Vector Laboratories, H-3300-250). Sections were then incubated with Alexa Fluor 647-conjugated streptavidin (Invitrogen, S21374) for 1 hour at room temperature. Nuclear counterstaining was performed by incubation with DAPI (300 nM) (Invitrogen, D1306) for 5 minutes at room temperature. Sections were mounted using Dako Fluorescence Mounting Medium (Agilent, S3023).

Fluorescence images were acquired using a BC43 benchtop confocal microscope (Oxford Instruments). Within each experiment, all images were collected using identical acquisition parameters to ensure consistency and enable qualitative comparison across tissues and conditions.

**Proteomic data processing and normalization**

Raw mass spectrometry data were processed using Proteome Discoverer (version 3.0) and searched against the UniProt mouse reference proteome. Peptide and protein identifications were filtered at a false discovery rate (FDR) of 1% using a target–decoy strategy. Unless otherwise indicated, only proteins identified with at least two unique peptides were retained for downstream analyses.

Detected proteins were mapped to UniProt “true proteins” (TP), defined as proteins annotated as secreted, cell membrane, or cell surface based on UniProt Subcellular Location (CC) annotations. When multiple protein groups mapped to the same gene symbol, the group with the highest number of unique peptides and the lowest missingness across samples was selected. Protein abundance was quantified using label-free quantification (LFQ) intensities and log₂-transformed after addition of a small pseudocount. For cross-sample comparisons, LFQ intensities were normalized and scaled using row-wise z-score transformation for visualization and clustering analyses. Missing values were handled in a context-dependent manner. For visualization and clustering analyses, missing values were left unfilled or set to zero after z-score transformation. For differential abundance and gene set–based analyses, missing values were imputed using minimal-value–based approaches to approximate low-abundance proteins while minimizing artificial inflation of fold changes.

**Dimensionality reduction and UMAP clustering**

To visualize global relationships among *in vivo*–labeled surface proteomes and assess tissue-level segregation, uniform manifold approximation and projection (UMAP) was applied to proteomic datasets following normalization and quality filtering.

For NHS-Biotin and HRP-mediated proximity labeling datasets, protein abundance matrices were restricted to UniProt-curated “true proteins”. Protein intensities were log₂-transformed and, where indicated, row-wise z-score normalization was applied. Missing values were handled as described above.

UMAP embeddings were computed using the uwot R package on scaled protein abundance matrices with cosine distance. Each biological replicate was treated as an independent observation. The number of nearest neighbors (n_neighbors) and minimum distance (min_dist) parameters were selected empirically for each dataset to optimize visualization of local structure and global tissue separation and are specified in the corresponding figure legends.

**Tissue-level proteome correlation analysis**

Pairwise correlation analysis was performed on tissue-averaged proteomic profiles derived from UniProt-annotated “true proteins” (TP) for both NHS-Biotin and CHRP datasets.

Pearson correlation coefficients were calculated using pairwise complete observations, and correlation matrices were visualized as heatmaps with hierarchical clustering applied to both rows and columns.

**Adipocyte-restricted HRP surfaceome profiling and dietary comparison**

Differential adipocyte surface proteomics analysis
For differential analysis of adipocyte surface proteins between chow and HFD conditions, protein intensity values were log₂-transformed after addition of a pseudocount of 1. Proteins were retained if detected in at least 2 of 3 biological replicates in both groups. Differential abundance was assessed using the limma R package by fitting a linear model to compare HFD versus chow samples. Log₂ fold change, nominal P values, and Benjamini–Hochberg adjusted P values were calculated for each protein. Differential abundance was visualized using volcano plots showing log₂ fold change and statistical significance. Selected candidate proteins were annotated using an exploratory threshold of nominal P < 0.05 and |log₂ fold change| > 0.5.

Gene set enrichment analysis
For comparative analysis of adipocyte surfaceomes under chow and HFD conditions, gene set enrichment analysis (GSEA) was performed using a pre-ranked approach. Proteins were ranked based on the difference in mean abundance between HFD and chow conditions across biological replicates. GSEA was conducted using the fgseaMultilevel algorithm implemented in the fgsea R package, with Hallmark gene sets from MSigDB obtained via the msigdbr package using mouse GeneSymbol identifiers. Gene sets were filtered to include pathways with sizes between 10 and 500 genes after intersection with the ranked protein list. Statistical significance was assessed using permutation-based testing, and pathways with FDR-adjusted P values below 0.05 were considered significant. Representative enriched pathways were selected for visualization based on statistical significance and normalized enrichment score.

Overlap analysis of chow and HFD adipocyte surfaceomes
Proteins were classified based on replicate-supported detection across diet conditions. Entries were collapsed to unique gene symbols, and protein presence within each group was defined as detection in at least 2 of 3 biological replicates. Proteins detected only in chow or HFD samples were designated chow-only or HFD-only, respectively, whereas proteins detected in both groups were classified as shared. Overlap between conditions was visualized using a Venn diagram.

**Deconvolution and subtype scoring**

To infer endothelial subtype patterns from CHRP proteomic profiles, we compared three complementary approaches using identical curated subtype marker sets from Supplementary Table S3: (1) z-score–based marker aggregation, (2) single-sample gene set enrichment analysis (ssGSEA), and (3) non-negative least squares (NNLS). For all methods, subtype scores were summarized at the tissue level and used for cross-method comparison.

For z-score–based marker aggregation, protein abundance values were standardized across samples by row-wise z-score transformation. Sample-level subtype scores were calculated as the mean z-scored abundance of detected marker proteins belonging to each subtype, followed by averaging across biological replicates within each tissue and mean-centering across tissues.

For ssGSEA-based scoring, subtype enrichment scores were computed using single-sample gene set enrichment analysis implemented in the GSVA R package. Within each sample, proteins were ranked by abundance, and enrichment scores were calculated based on the relative distribution of subtype marker proteins within the ranked list compared to the background proteome. Scores were subsequently standardized by row-wise z-score transformation and averaged across biological replicates to generate tissue-level subtype enrichment profiles.

For NNLS-based deconvolution, a gene-by-subtype signature matrix was constructed from detected marker genes as a binary indicator matrix normalized by marker set size. Tissue-level proteomic profiles were obtained by averaging protein abundance across biological replicates. Subtype coefficients were estimated by modeling each tissue-level proteomic vector as a non-negative linear combination of subtype signatures. Coefficients were normalized within each tissue and mean-centered across tissues for comparison.

Agreement across methods was assessed using pairwise Pearson correlation of tissue-level subtype score matrices. For each endothelial subtype, correlations were calculated across tissues for z-score–based aggregation versus ssGSEA, z-score–based aggregation versus NNLS, and ssGSEA versus NNLS, and visualized as a heatmap. In addition, relationships among endothelial subtypes were evaluated using Spearman correlation to capture monotonic associations independent of scale.

**Cross-tissue comparison of endothelial membrane proteins detected by CHRP proteomics and scRNA-seq**

To systematically compare endothelial membrane proteins detected by *in vivo* CHRP-based surface proteomics and single-cell RNA sequencing (scRNA-seq), we implemented a cross-tissue, modality-aware analysis framework across six tissues. Endothelial transcriptomic references were derived from publicly available single-cell RNA sequencing datasets, including the Murine Endothelial Cell Atlas (E-MTAB-8077; Kalucka et al.) and adipose tissue datasets GSM5359340, GSM5359345, and GSM5359346. For non-adipose tissues, endothelial-specific expression metrics were computed from EC-resolved count matrices derived from the Kalucka atlas, including mean expression (meanLogCP10K) and detection rate across endothelial cells. For adipose tissue, endothelial expression profiles were reconstructed from raw data using a standardized Seurat workflow, followed by endothelial cell identification and calculation of EC specificity metrics.

For each tissue, membrane proteins identified by CHRP proteomics (gene symbols) were compared with endothelial-expressed genes derived from tissue-matched scRNA-seq references. Proteins were classified into three non-overlapping sets: CHRP-only (detected by proteomics but not scRNA-seq), scRNA-seq–only (sc-only; detected by scRNA-seq but not proteomics), and overlap (detected by both modalities). For cross-tissue analyses, modality-specific gene sets were additionally aggregated using the union of detected genes across tissues within each modality.

Membrane proteins were assigned to 11 functional classes based on curated Gene Ontology Cellular Component annotations. Class composition was summarized using both pooled aggregation across tissues and tissue-resolved analyses. Tissue-specific enrichment or depletion of each functional class in CHRP-only or sc-only sets relative to the overlap set was quantified using log₂ odds ratios derived from Fisher’s exact tests. Association strength between functional class and detection modality within each tissue was summarized using Cramér’s V based on class-by-modality contingency tables.

For modality-specific detection metrics, CHRP signal intensity was defined as the log₂-transformed mean protein abundance per gene. For scRNA-seq–derived metrics, detection rate was defined as the fraction of endothelial cells expressing a given gene and summarized as the log₁₀-transformed detection rate. Detection breadth was defined as the log₂ fold-change of the number of endothelial cells expressing each gene relative to the median value of scRNA-seq–only genes within the same tissue, with a pseudocount of 1 added to both numerator and denominator to stabilize low-count estimates.

Differences between protein sets were assessed using Wilcoxon rank-sum tests, with effect size summarized by group medians and P values adjusted using the Benjamini–Hochberg method. For pooled cross-tissue analyses, detection metrics were normalized within each tissue prior to aggregation.

**Metabolic membrane protein analysis**

Metabolic membrane proteins were defined from CHRP-detected proteins using a strict membrane-based annotation strategy. Proteins were first filtered based on UniProt subcellular localization annotations to retain membrane-associated proteins, including proteins annotated as cell membrane, plasma membrane, transmembrane, integral membrane, GPI-anchored, or membrane raft-associated proteins. Proteins annotated exclusively as secreted, extracellular matrix, or extracellular region proteins without explicit membrane annotation were excluded. Mitochondrial proteins, including mitochondrial membrane proteins, were also excluded. The resulting strict membrane protein set was intersected with MSigDB metabolism-associated gene sets to define metabolic membrane proteins for downstream analyses.

Protein abundance values were averaged across biological replicates within each tissue. For visualization, tissue-averaged abundance values were row-scaled across tissues to generate z-score matrices. Dimensionality reduction and correlation analyses were performed using the strict membrane metabolic protein matrix. UMAP was performed on replicate-level scaled abundance values using cosine distance, and tissue-level correlation analysis was performed using Pearson correlation of tissue-averaged scaled abundance profiles.

To identify tissue-enriched metabolic membrane proteins, each protein was assigned to the tissue in which it showed the highest mean abundance, defined as the primary tissue. A tissue enrichment ratio was calculated relative to the second-highest tissue and further filtered to retain proteins with primary-tissue mean log₂ abundance ≥3, detection in at least 2 of 3 biological replicates in the primary tissue, and enrichment ratio ≥1.2 relative to the second-highest tissue. Proteins dominated by vesicle trafficking, endocytosis, generic signaling adaptor, cytoskeletal, were excluded from the visualization set. Candidates were ranked within each primary tissue using a composite score incorporating tissue enrichment, primary-tissue abundance, and replicate support. Up to the top six candidates per tissue were selected for heatmap visualization.

For module-level analysis, metabolic membrane proteins were grouped into curated functional modules based on biological function and pathway annotation, including glucose transport, amino acid transport, lactate transport, lipid metabolism, and purinergic signaling. Module scores were calculated as the mean row-scaled abundance of detected proteins within each module.

Gene Ontology (GO) enrichment analysis was performed using tissue-assigned metabolic membrane protein sets. Genes were converted to Entrez identifiers and GO biological process enrichment was performed using clusterProfiler with Benjamini-Hochberg FDR correction. GO terms related to metabolism, metabolite transport, lipid handling, solute transport, redox-related processes, and extracellular metabolic signaling were retained for visualization. Enriched terms were displayed using bubble plots, with point size indicating gene count and color indicating −log₁₀(FDR).

**Identification and annotation of tissue-selective endothelial membrane protein candidates**

Protein abundances were log₂-transformed and averaged across biological replicates for each tissue. For each protein, the tissue with the highest mean abundance was defined as the primary tissue, and tissue enrichment was calculated as the difference between the primary-tissue mean log₂ abundance and the second-highest tissue mean log₂ abundance (tissue_diff). Candidate proteins were restricted to UniProt-annotated membrane-associated proteins, including cell membrane, plasma membrane, transmembrane, integral membrane, GPI-anchored, or membrane raft annotations. Proteins annotated only as secreted, extracellular matrix, extracellular space, or basement membrane proteins, as well as mitochondrial proteins, were excluded. Proteins were retained as tissue-selective endothelial membrane candidates if they had tissue_diff ≥1, primary tissue mean log₂ intensity ≥3, and detection in at least 2 of 3 biological replicates in the primary tissue. The complete candidate list is provided in Table S6.

For heatmap visualization, candidates were ranked within each primary tissue using a composite proteomics-based priority score: 0.50 × normalized tissue enrichment + 0.35 × normalized primary-tissue abundance + 0.15 × replicate support. Replicate support was defined as the fraction of detected biological replicates in the primary tissue. This score was used only for visualization and did not affect inclusion in Table S6. The top 30 candidates per tissue were displayed using row-scaled z-scores.

Cross-tissue distribution was assessed using the full membrane candidate table. Tissue membership was assigned when tissue-level mean log₂ abundance was ≥20, and candidates were classified as tissue-unique, shared across 2-3 tissues, or shared across ≥4 tissues. Exact tissue intersections were calculated for all combinations, and the 15 most frequent intersections were shown in the UpSet-like plot.

For compound-target annotation, candidates were cross-referenced with ChEMBL 30 database. The ChEMBL 30 SQLite database was downloaded from the ChEMBL release archive (<http://ftp.ebi.ac.uk/pub/databases/chembl/ChEMBLdb/releases/chembl_30>). Candidates were considered ChEMBL-matched if linked to at least one ChEMBL compound through drug-mechanism annotation. For alluvial visualization, up to five ChEMBL-matched candidate proteins per tissue and up to two compounds per protein were displayed. The complete compound-target annotation output is provided in Table S7.

GO molecular function enrichment was performed using clusterProfiler with org.Mm.eg.db. Tissue-selective membrane candidates from each primary tissue were used as foreground, and all CHRP-detected membrane proteins were used as background. Enrichment was assessed using Benjamini-Hochberg FDR-adjusted P values, with p <0.05, q <0.20, and GO gene-set size 10-500. Dotplots were generated for tissues with significant enriched terms.

**Statistical analysis**

Statistical analyses were performed using R (version ≥4.2.0) unless otherwise specified. No statistical methods were used to predetermine sample size. Biological replicates represent independent mice or independently processed tissue samples, as specified in the figure legends.

For quantitative proteomic analyses, protein abundance values were log₂-transformed prior to statistical testing. Statistical analyses were performed using methods appropriate for each dataset, as described in the corresponding Methods sections. Multiple hypothesis testing was controlled using the Benjamini–Hochberg false discovery rate (FDR) procedure, with significance thresholds specified for each analysis.

Correlation analyses were performed using Pearson or Spearman correlation coefficients, as indicated in the corresponding figure legends. Hierarchical clustering was conducted using standard distance metrics and linkage methods as described in the relevant Methods sections.

All statistical tests were two-sided. Exact statistical tests, sample sizes, and significance thresholds are reported in the figure legends. No data were excluded unless explicitly stated. All software used is publicly available or commercially licensed and is listed in the Key Resources Table.

1.	Wang, Y., et al., *Ephrin-B2 controls VEGF-induced angiogenesis and lymphangiogenesis.* Nature, 2010. **465**(7297): p. 483-6.

**Key Resources Table**

| **Chemicals, Reagents, Media and Kits** | **SOURCE** | **IDENTIFIER** |
| --- | --- | --- |
| Sulfo-NHS-LC-Biotin | ApexBio Technology | A8003 |
| Biotin-XX Tyramide | ApexBio Technology | A8012 |
| 60% high-fat diet | BioServ | S1850 |
| Tamoxifen | Sigma-Aldrich | T5648-1G |
| Glycine | Sigma-Aldrich | G7126-500G |
| Sodium chloride | Sigma-Aldrich | S7653-1KG |
| Calcium chloride hexahydrate | Sigma-Aldrich | 442909-1KG |
| Potassium Chloride (3 M) | Boston Bioproducts | MT-253 |
| Magnesium chloride solution | Sigma-Aldrich | M1028-100ML |
| Sodium phosphate monobasic dihydrate | Sigma-Aldrich | 71500-1KG |
| Glucose | Sigma-Aldrich | D9434-500G |
| HEPES Buffer (1 M, pH 7.4) | Boston Bioproducts | BBH-74 |
| Hydrogen peroxide solution | Sigma-Aldrich | 216763-100ML |
| Sodium azide | Sigma-Aldrich | S2002-5G |
| (+)-Sodium L-ascorbate | Sigma-Aldrich | A7631-25G |
| (±)-6-Hydroxy-2,5,7,8-tetramethylchromane-2-carboxylic acid | Sigma-Aldrich | 238813-1G |
| UltraPure™ 1M Tris-HCI, pH 8.0 | Invitrogen | 15568025 |
| Sodium Dodecyl Sulfate, SDS (20%) | Boston Bioproducts | BM-230 |
| Sodium deoxycholate | Thermo Scientific | 89904 |
| Triton™ X-100 | Sigma-Aldrich | X100-500ML |
| Halt™ Protease and Phosphatase Inhibitor Cocktail (100X) | Thermo Scientific | 78442 |
| PMSF: Phenylmethylsulfonyl fluoride | Thermo Scientific | 36978 |
| Streptavidin magnetic beads | Thermo Scientific | 88817 |
| Sodium carbonate | Sigma-Aldrich | S2127-1KG |
| Urea | Sigma-Aldrich | U5378-100G |
| Pierce™ Lane Marker Non-Reducing Sample Buffer | Thermo Scientific | 39001 |
| 2-Mercaptoethanol | Bio-Rad | 1610710 |
| Odyssey One-Color Protein Molecular Weight Marker | Li-Cor | 92840000 |
| D-(+)-Biotin (CAS 58-85-5) | Santa Cruz | sc-204706B |
| DAPI | Invitrogen | D1306 |
| Antigen Unmasking Solution, Citric Acid Based | Vector Laboratories | H-3300-250 |
| Dako Fluorescence Mounting Medium | Agilent | S3023 |
| Pierce™ BCA Protein Assay Kits | Thermo Scientific | 23227 |
| Pierce™ Silver Stain for Mass Spectrometry | Thermo Scientific | 24600 |

| **Software and Packages** | **SOURCE** | **Version** |
| --- | --- | --- |
| Rstudio IDE | Rstudio | 2024.12.1.563 |
| R | R Foundation for Statistical Computing | 4.5.0 |
| readxl | CRAN | 1.4.5 |
| dplyr | CRAN | 1.1.4 |
| tibble | CRAN | 3.3.0 |
| stringr | CRAN | 1.5.2 |
| ggplot2 | CRAN | 4.0.0 |
| scales | CRAN | 1.4.0 |
| uwot | CRAN | 0.2.3 |
| readr | CRAN | 2.1.5 |
| ggforce | CRAN | 0.5.0 |
| Seurat | CRAN | 5.3.0 |
| tidyr | CRAN | 1.3.1 |
| SeuratObject | CRAN | 5.2.0 |
| BiocManager | CRAN | 3.22 |
| pheatmap | CRAN | 1.0.13 |
| RColorBrewer | CRAN | 1.1.3 |
| svglite | CRAN | 2.2.2 |
| GSVA | Bioconductor | 2.2.1 |
| msigdbr | CRAN | 25.1.1 |
| ComplexHeatmap | Bioconductor | 2.24.1 |
| grid | R (base) | R 4.5.0 |
| circlize | CRAN | 0.4.16 |
| ggrepel | CRAN | 0.9.6 |
| purrr | CRAN | 1.1.0 |
| dendextend | CRAN | 1.19.1 |
| GSEABase | Bioconductor | 1.70.1 |
| limma | Bioconductor | 3.64.3 |
| VennDiagram | CRAN | 1.7.3 |
| clusterProfiler | Bioconductor | 4.16.0 |
| org.Mm.eg.db | Bioconductor | 3.21.0 |
| ontologyIndex | CRAN | 2.12 |
| AnnotationDbi | Bioconductor | 1.70.0 |
| GO.db | Bioconductor | 3.21.0 |
| openxlsx | CRAN | 4.2.8 |
| fgsea | Bioconductor | 1.34.2 |
| forcats | CRAN | 1.0.1 |
| Matrix | CRAN | 1.7.4 |
| data.table | CRAN | 1.17.8 |
| patchwork | CRAN | 1.3.2 |
| enrichplot | Bioconductor | 1.28.4 |
| ComplexUpset | CRAN | 1.3.6 |
| cowplot | CRAN | 1.2.0 |
| qvalue | Bioconductor | 2.40.0 |
| withr | CRAN | 3.0.2 |
| matrixStats | CRAN | 1.5.0 |
| pcaMethods | Bioconductor | 2.0.0 |
| imputeLCMD | CRAN | 2.1 |
| glmnet | CRAN | 4.1.10 |
| impute | Bioconductor | 1.82.0 |
| writexl | CRAN | 1.5.4 |
| tools | R (base) | 4.5.0 |
| edgeR | Bioconductor | 4.6.3 |
| FNN | CRAN | 1.1.4.1 |
| DOSE | Bioconductor | 4.2.0 |
| DBI | CRAN | 1.2.3 |
| RSQLite | CRAN | 2.4.3 |
| ggalluvial | CRAN | 0.12.5 |
